# Lineage–specific amino acids define functional attributes of the protomer-protomer interfaces for the Rad51 and Dmc1 recombinases

**DOI:** 10.1101/2024.12.03.626531

**Authors:** Mike Petassi, Yeonoh Shin, Aidan M. Jessop, Katherine Morse, Stefan Y. Kim, Razvan Matei, Vivek B. Raina, Eric C. Greene

**Author notes:** Equal contribution (Mike Petassi, Yeonoh Shin).

## Abstract

Most eukaryotes possess two Rad51/RecA family DNA recombinases that are thought to have arisen from an ancient gene duplication event: Rad51, which is expressed in both mitosis and meiosis; and Dmc1, which is only expressed in meiosis. To explore the evolutionary relationship between these recombinases, here, we present high-resolution CryoEM structures of *S. cerevisiae* Rad51 filaments and *S. cerevisiae* Dmc1 filaments bound to ssDNA, which reveal a pair of stacked interfacial aromatic amino acid residues that are nearly universally conserved in Rad51 but are absent from Dmc1. We use a combination of bioinformatics, genetic analysis of natural sequence variation, and deep mutational analysis to probe the functionally tolerated sequence space for these stacked aromatic residues. Our findings demonstrate that the functional landscape of the interfacial aromatic residues within the Rad51 filament is highly constrained. In contrast, the amino acids at the equivalent positions within the Dmc1 filament exhibit a broad functional landscape. This work helps highlight the distinct evolutionary trajectories of these two eukaryotic recombinases, which may have contributed to their functional and mechanistic divergence.

**AUTHOR SUMMARY:** Most eukaryotic organisms have two closely related proteins, Rad51 and Dmc1, that are needed for different aspects of genetic recombination. These proteins may have evolved from a single gene that was duplicated during the early evolution of eukaryotes. Rad51 is active during both normal cell division (mitosis) and sexual reproduction (meiosis), while Dmc1 is only active during meiosis. To better understand how these proteins are related, we studied their three-dimensional structures using high resolution cryogenic electron microscopy. Our findings show that Rad51 has a specific set of conserved amino acids located at the protein interfaces, but this set of amino acids is different in Dmc1. We used a series of genetic approaches to analyze how these amino acids affect the proteins’ function. Our results show that Rad51 has a strict set of rules governing the identify of these amino acids, whereas Dmc1 does not. This research sheds light on how Rad51 and Dmc1 have evolved differently, leading to distinct functions in genetic recombination.

## INTRODUCTION

Homologous recombination (HR) contributes to the maintenance of genome integrity among all kingdoms of life and serves as a major driving force in evolution (1, 2). During HR, a presynaptic single–stranded DNA (ssDNA) is paired with the complementary strand of a homologous double–stranded DNA (dsDNA), resulting in displacement of the non–complementary strand (3–6), and the resulting displacement loop (D–loop) intermediates can then be channeled through several mechanistically distinct pathways to complete repair (3–6). HR plays roles in double-strand DNA break (DSB) repair (6, 7), the rescue of stalled or collapsed replication forks (8, 9), chromosomal rearrangements (10–12), horizontal gene transfer (13), and meiosis (14–16).

The protein participants, nucleoprotein structures, and general HR reaction mechanisms are broadly conserved (3–5, 17). The DNA pairing reactions that take place during HR are promoted by the Rad51/RecA family of DNA recombinases, which are ATP–dependent proteins that form extended helical filaments on single stranded DNA, that are referred to as presynaptic complexes (3–5, 17). Crystal structures of RecA–ssDNA presynaptic and RecA–dsDNA postsynaptic complexes reveal that the DNA is organized into near B–form base triplets separated by ∼8 Å between adjacent triplets (18). This structural organization likely underpins homology recognition and the ability of the Rad51/RecA family of recombinases to promote DNA strand invasion in 3–nt steps (18–22).

Interestingly, most eukaryotes have two Rad51/RecA family recombinases: Rad51, which is constitutively expressed; and Dmc1, which is only expressed during meiosis (14, 16, 23–25). Rad51 and Dmc1 arose from an ancient gene duplication event, and the emergence of Dmc1 as a separate lineage may have coincided with the emergence of meiosis and sexual reproduction (26–29). These two proteins remain closely related; for instance, *S. cerevisiae* Rad51 and Dmc1 share 45% sequence identity and 56% sequence similarity and both proteins perform the same basic biochemical function, namely the pairing of homologous DNA sequences (14, 16). While Rad51 and Dmc1 do share ∼45% amino sequence identity across species, they also harbor amino acids that are conserved within either the Rad51 lineage, or the Dmc1 lineage, but not both (27, 30). The functional significance of these lineage-specific amino acids remains largely unexplored.

Rad51 and Dmc1 were identified over 25 years ago (23, 31), yet we still have a poor understanding of why most eukaryotes require these two recombinases (14, 16). Prevailing hypotheses are that (*i*) each recombinase is required to interact with a specific subset of mitotic- or meiotic-specific accessory factors, (*ii*) there are biochemical differences between the recombinases making each uniquely suited to their roles in mitotic or meiotic HR, or (*iii*) both (14, 16). We favor the latter hypothesis, given that there are examples of Rad51- and Dmc1-specific interacting factors (discussed below)(14, 16), and prior work revealed that Rad51 and Dmc1 respond differently when presented with mismatch–containing HR intermediates (30, 32, 33). Nevertheless, it remains unclear why eukaryotes have evolved two recombinases (14, 16).

To help further understand the evolutionary relationship between Rad51 and Dmc1, we have begun addressing the question of whether differences in protein sequence and structure may underly differences in biological function. Here we present a side-by-side comparison of the CryoEM structures of ssDNA–bound nucleoprotein filaments of these two recombinases from *Saccharomyces cerevisiae*. As part of this comparison, we analyze a pair of stacked tyrosine residues located at the interface between adjacent *S. cerevisiae* Rad51 monomers within the nucleoprotein filaments. Bioinformatic analysis reveals that stacked aromatic residues at these positions are highly conserved within the Rad51 lineage of the Rad51/RecA family of recombinases. In contrast, the stacked aromatic residues are absent from the Dmc1 lineage of the Rad51/RecA family of recombinases. For example, the two tyrosine residues found in *S. cerevisiae* Rad51 are instead replaced with serine and leucine in *S. cerevisiae* Dmc1. Using a combination of genetic assays with natural sequence variants and deep mutagenic screens to test all possible combinations of amino acid residues we show that Rad51 has very strict requirements for these interfacial residues, whereas Dmc1 exhibits a striking degree of structural plasticity. Our work helps to shed further light on the unique attributes of the Rad51 and Dmc1 lineages of the Rad51/RecA family of recombinases.

## RESULTS

### Rad51– and Dmc1–ssDNA filament structures

We solved the CryoEM structure of *S. cerevisiae* Rad51 bound to a 96–nt ssDNA substrate to 2.7 Å resolution (Fig. 1a-c, Fig. S1a-c, Table S1). The Rad51 structure we have obtained is similar to the previous 3.25 Å crystal structure of yeast Rad51 (34), but with a few important differences. First, the crystal structure used an N-terminally truncated (amino acids 1-79) version of Rad51–I345T, which is a gain-of-function mutant with enhanced ssDNA binding activity (35), whereas our structure was solved using full-length wild-type Rad51. Second, the crystal structure was solved with the non-hydrolysable ATP analog ATPγS, but the nucleotide analog was not observed in the structure, instead only a sulfate ion was visible in the nucleotide binding pocket (34). Our Rad51 structure was solved in the presence of ATP, and the entire ATP molecule is visible within the nucleotide-binding cleft (Fig. a-b). Third, the ssDNA used for crystallography was not visible within the Rad51 filament structure (34), whereas the ssDNA molecule is visible within our CryoEM structure (Fig. 1a-b). Fourth, our CryoEM structure shows a 105 Å pitch with 6.46 Rad51 monomers per turn, which contrasts with the more elongated 130 Å pitch observed in the crystal structure (34). Fifth, the L1 and L2 DNA binding loops were not visible in the crystal structure (34), whereas we can resolve the entirety of L1 and a portion of L2 (L2 residues D332 to D340 cannot be assigned due to poor density), including the amino acid residues that contact the ssDNA (Fig. 1a). We also solved a CryoEM structure of *S. cerevisiae* Dmc1 bound to a 96–nt ssDNA substrate to 2.6 Å resolution and the structure shows a pitch of 98.4 Å with 6.35 Dmc1 monomers per turn (Fig. 1d-f, Fig. S1d-f, Table S1). As with Rad51, our structure for the Dmc1 filament utilized the full-length wild-type protein, the complex is in the ATP-bound state, and both the ATP and ssDNA are visible within the structure. Note also that a 3.2 Å Dmc1 CryoEM structure was recently reported and agrees with our structure (32).

**Figure 1.**
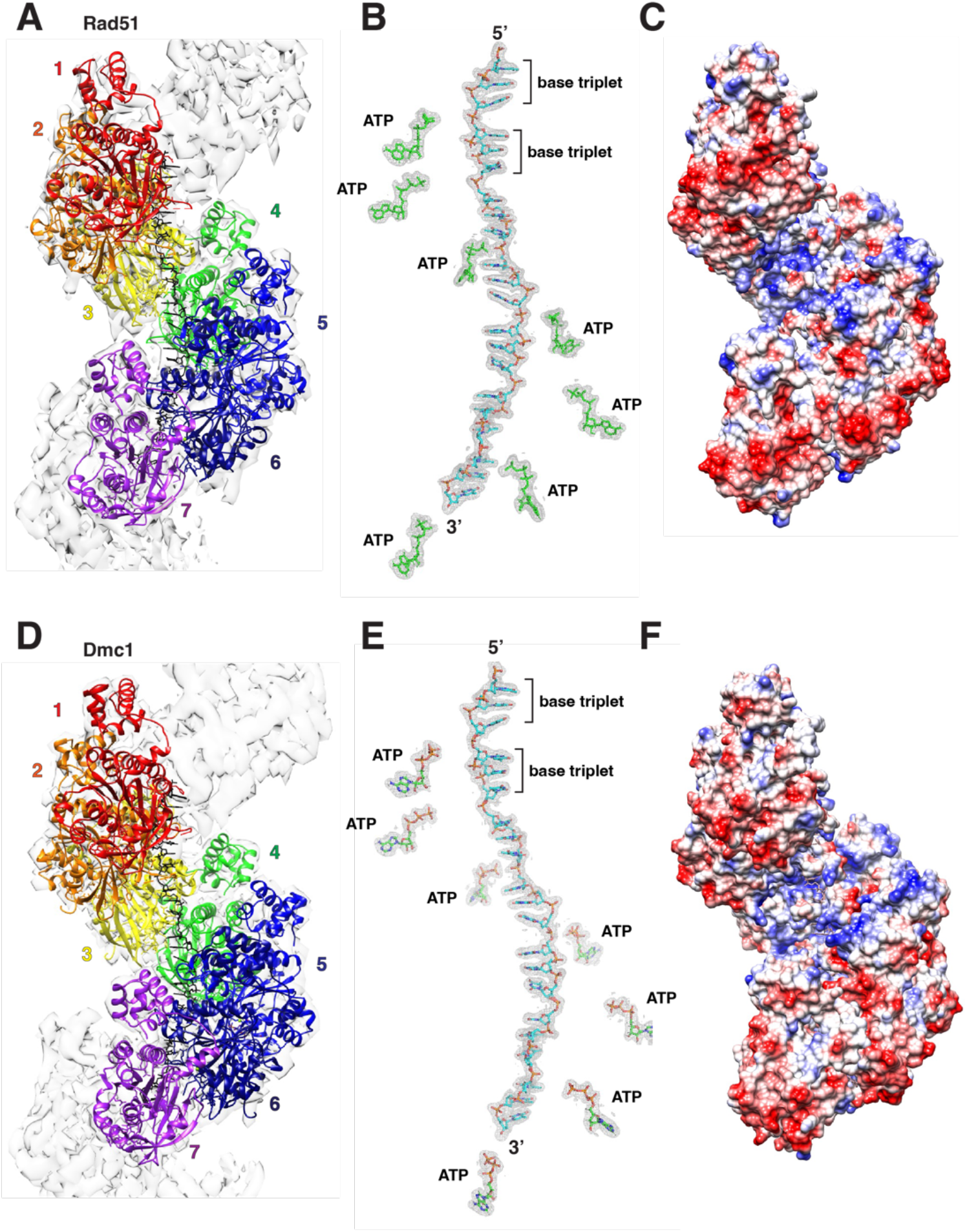
Structures of the *S. cerevisiae* Rad51 and Dmc1 filaments. a, CryoEM structure of yeast Rad51 bound to an ssDNA fragment in the presence of ATP. Each Rad51 monomer within the filament is highlighted in a different color. **b,** Structure of the ssDNA from within the Rad51 filament (proteins are not shown) and location of the bound ATP molecules. **c,** Electrostatic surface diagram of the Rad51 filament. **d,** CryoEM structure of yeast Dmc1 bound to an ssDNA fragment in the presence of ATP. Each Dmc1 monomer within the filament is highlighted in a different color. **e,** Structure of the ssDNA from within the Dmc1 filament (proteins are not shown) and location of the bound ATP molecules. **f,** Electrostatic surface diagram of the Dmc1 filament.

### Comparison of Rad51–ssDNA and Dmc1–ssDNA monomers

Consistent with their high degree of sequence conservation, the structures of the Rad51– and Dmc1–ssDNA filaments were highly similar and an overlay of the Rad51 and Dmc1 monomers revealed an RSMD of just 1.05 Å (Fig. 2a-c). Both proteins have a highly conserved core domain, which contains the ATP-binding residues on the 3’ monomer face relative to the orientation of the ssDNA (Fig. 2a-c). The ssDNA is held in place by the two DNA-binding loops positioned near either end of the base triplet with L1 positioned near the 5’ end of the base triplet and L2 positioned at the 3’ near end of the base triplet (Fig. 2a-c). Both proteins have an N-terminal domain that extends from the globular core and is comprised of four helix bundle that folds back to contact the core domain (Fig. 2a- c). All of the features seen here are largely consistent with previously reported structural data (32, 34) and further reinforce the close relationship between Rad51 and Dmc1.

**Figure 2.**
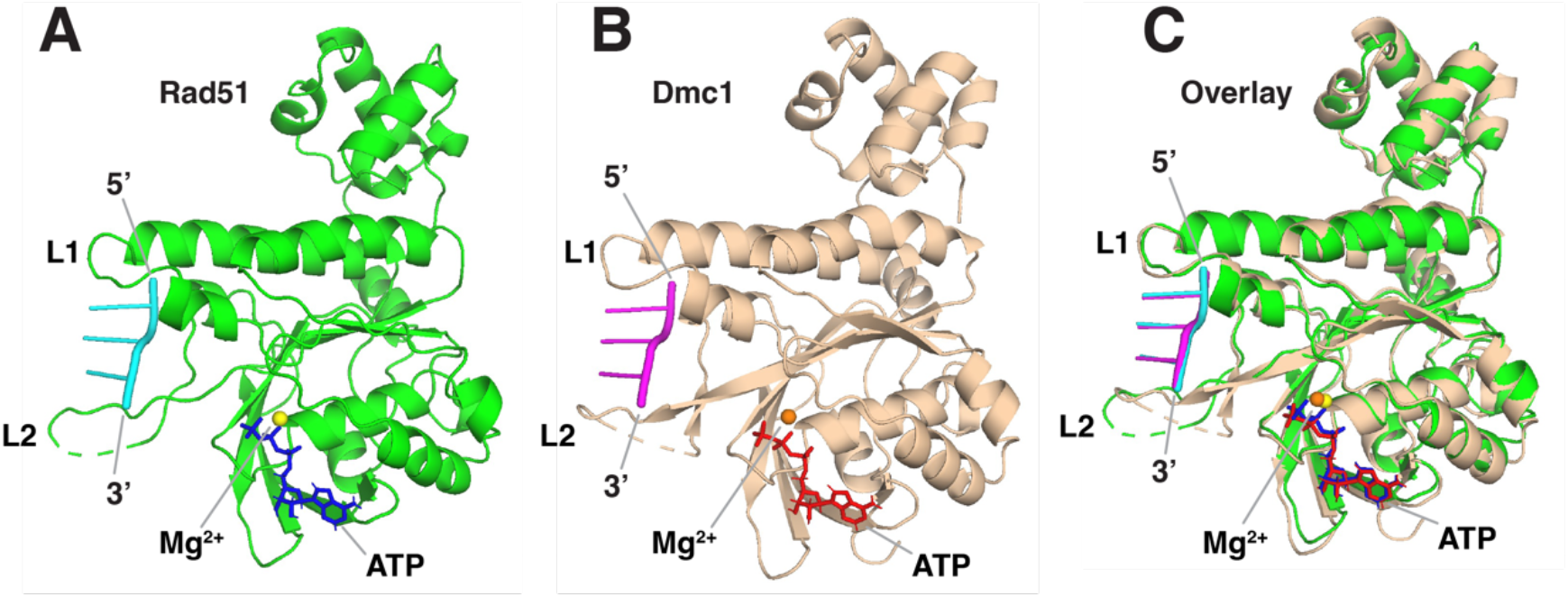
Comparison of Rad51 and Dmc1 monomers. a, Structure of a single Rad51 protein monomer (shown in green) from within the Rad51 filament including the bound ssDNA (shown in cyan) and ATP molecule (shown in blue). **b,** Structure of a single Dmc1 protein monomer (tan) from within the Dmc1 filament including the bound ssDNA (shown in magenta) and ATP molecule (shown in red). **c,** Overlay of the Rad51 and Dmc1 monomer structures.

### Unique protomer-protomer interfaces for Rad51 and Dmc1 filaments

Our structure of Rad51 highlights a pair of stacked tyrosine (Y) residues within the Rad51-Rad51 interface (Y112 & Y253; Fig. 3a-d), which is consistent with prior structures of yeast and human Rad51 (34, 36). Remarkably, we find that this YY stacking interaction is completely absent from Dmc1 (Fig. 3a-d). Instead, in *S. cerevisiae* Dmc1 the corresponding residues are S48 and L189 (Fig. 3d). Bioinformatic analysis reveals that the stacking interaction is highly conserved within the Rad51 lineage, with 93.9% of available sequences (N = 1,186 Eukarya) being comprised of either YY (46.8%), YF (35.7%) or FY (11.5%) pairs (Fig. 3e-f; Supplementary data file). In Dmc1, these residues are most often MY (55.5%), GL (10.6%) and SL (9.3%); we have identified only two examples of stacked aromatic residues for Dmc1, both are species of parasitic plant fungi from the genus *Coleophoma* (Fig. 3g-h; Supplementary data file). Thus, the stacking interaction at the Rad51 monomer-monomer interface appears to be uniquely conserved within the Rad51 lineage of the Rad51/RecA family of recombinases and is not conserved within the Dmc1 lineage.

**Figure 3.**
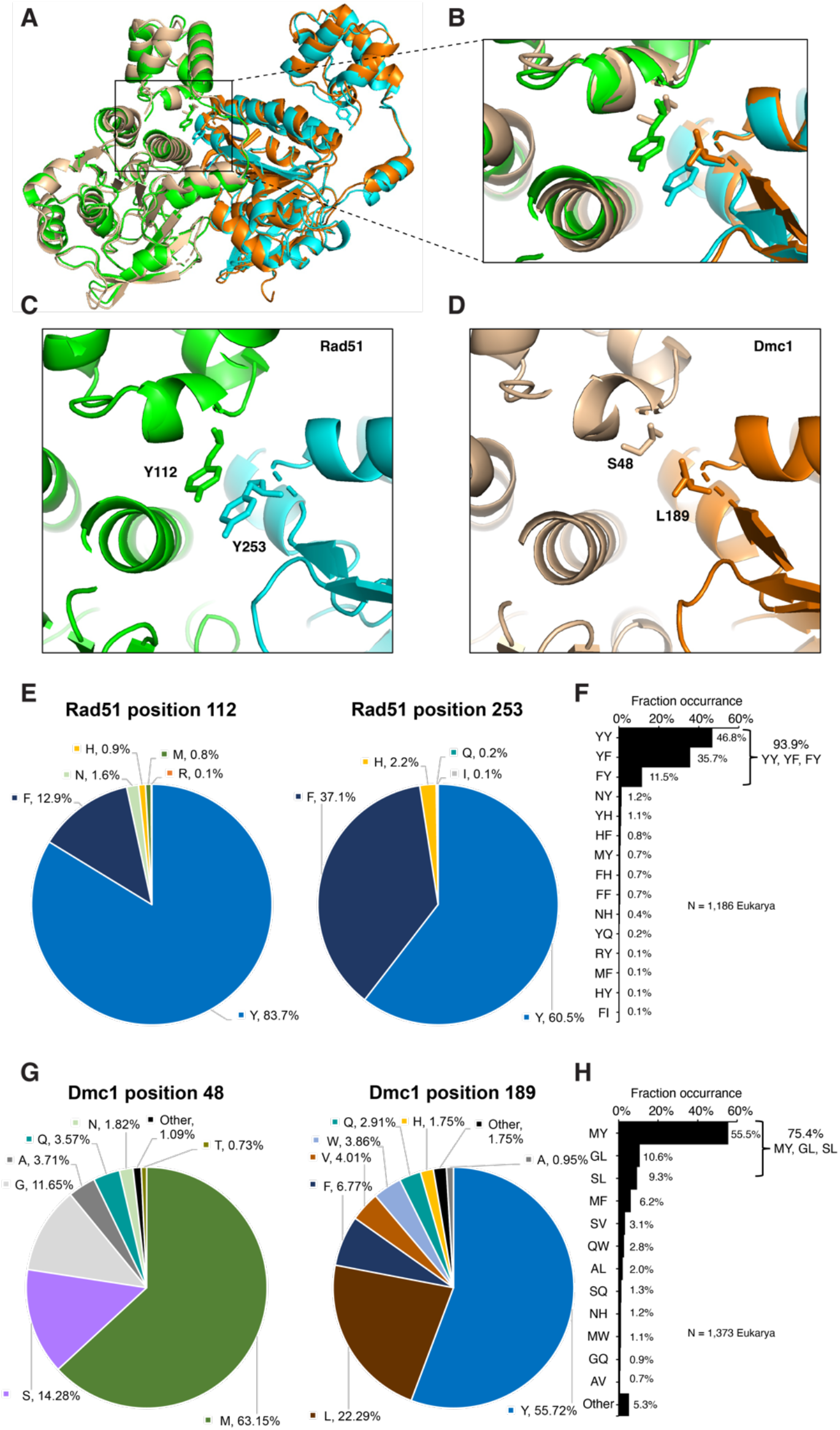
Stacked aromatic residues are a conserved feature of the Rad51 monomer- monomer interface. a, An overlay of two adjacent Rad51 (green and cyan) and Dmc1 (tan and orange) monomers from with each respective nucleoprotein filament. The side chains for Rad51 residues Y112 and 253, and Dmc1 residues S48 and L189 are shown. **b,** Overlay highlight the Rad51 and Dmc1 protein-protein interfaces. **c,** Structure of the Rad51 protein-protein interface showing the locations of the stacked tyrosine residues (Y112 and Y253). **d,** Structure of the Dmc1 protein-protein interface showing the locations of the residues S48 and L189. **e,** Amino acid residue conservation at positions equivalent to *S. cerevisiae* Rad51 Y112 and Y253. **f,** Frequency of naturally occurring Rad51 interfacial amino acid residue pairs (see Supplementary data file). **g,** Amino acid residue conservation at positions equivalent to *S. cerevisiae* Dmc1 S48 and L189. **h,** Frequency of naturally occurring Dmc1 interfacial amino acid residue pairs (see Supplementary data file).

Further bioinformatic analysis of the natural Rad51 sequence variants shows that there is just a limited subset of amino acid residues (8 in total; Y, F, H, N, M, Q, R, and I) found at the positions equivalent to *S. cerevisiae* Y112 and Y253, and consists primarily of Y and F (*i.e.* Y, 1710 examples; F, 595 examples; H, 37 examples; N, 19 examples; M, 9 examples; Q, 2 examples; R, 1 example; I, 1 example)(Fig. 3f & Supplementary data file); note that we do not rule out the possibility that the rarer amino acid residues (Q, R, I) may reflect sequencing errors. The remaining 12 amino acid residues (K, D, E, S, T, C, G, A, V, L, P, and W) are all excluded from our data set of natural Rad51 sequence variants at the positions equivalent to *S. cerevisiae* Y112 and Y253 (Fig. 3e-f). In striking contrast, there appear to be no examples of amino acid residues that are completely excluded from these positions in Dmc1 (Fig. 3g-h & Supplementary data file), although some of the amino acids are very rare (*i.e.* C, 2 examples; D, three examples; E, 1 example; I, 4 examples; K, 1 example; P, 2 examples; R, 1 example); again we do not rule out the possibility that these rare amino acid residues may reflect database sequencing errors. Regardless, our analysis of the natural Rad51 and Dmc1 sequence variants further suggest that the functional amino acid landscape at the YY positions in Rad51 is highly restricted, whereas that of Dmc1 may be more flexible.

### Importance of the stacked aromatics for Rad51 activity

Our bioinformatic analysis suggests that stacked aromatic residues at positions corresponding to *S. cerevisiae* Rad51 Y112 and Y253 are highly conserved throughout the Rad51 lineage, raising the question of whether these amino acid residues are important for Rad51 function. To test the importance of these residues, we generated mutants in which we replace Y112 and Y253 with alternative sets of amino acid residues. For the first set of mutants, the stacked tyrosine residues were changed to glutamate (Y112E) and lysine (Y253K), either individually or in combination; we reasoned that if a physical interaction was necessary for Rad51 function, then it may be possible to functionally replace the stacked tyrosines with a salt bridge. We also generated a mutant in which we replaced the tyrosine residues from Rad51 with the serine (Y112S) and leucine (Y253L) residues from *S. cerevisiae* Dmc1. These mutants were then tested for function in genetic, biochemical, and single molecule assays.

Genes encoding these alleles were integrated into chromosome V under the control of the endogenous *RAD51* promoter and the resulting *S. cerevisiae* W303 strains were assayed for growth in the presence of the DNA-alkylating agent methyl methane sulfonate (MMS). None of the *rad51* mutants were able to support cell growth on plates with MMS, suggesting that these proteins may have been functionally inactive (Fig. S2a). In addition, none of the mutants were able to support cell survival even when overexpressed (Fig. S2b). Lastly, none of the mutant proteins were capable of supporting D-loop formation in bulk biochemical assays (Fig. S2c-d) and none were able to assemble into nucleoprotein filaments on single-stranded DNA in single molecule DNA curtain assays (Fig. S2e-f). Taken together, our data suggests the YY stacking interaction is necessary for Rad51 biological function and replacing these residues with either a salt bridge (Y112E, Y253K) or the equivalent amino acid residues from Dmc1 (Y112S, Y253L) results in inactive proteins that are defective for nucleoprotein filament assembly.

### Functional analysis of Rad51 natural sequence variants

We sought to further assess the functional constraints on the stacked aromatic residues found at the Rad51 protomer interfaces by asking whether the naturally occurring sequences variants found in our bioinformatic analysis could retain function when placed within the context of *S. cerevisiae* Rad51. For this, we constructed a total of 14 sequence variants, which together with the stacked YY pair found in *S. cerevisiae*, account for all of the natural amino sequence variants identified in our bioinformatic analysis. Genes encoding these alleles were integrated into chromosome V under the control of the endogenous *RAD51* promoter (Fig. 4a) and the resulting strains were assayed for growth in the presence of MMS (Fig. 4b). Control assays confirmed that the wild-type *RAD51* strain grew under at all concentrations of MMS tested, whereas the negative control *rad51τι* strain did not survive at any MMS concentration tested (Fig. 4b).

**Figure 4.**
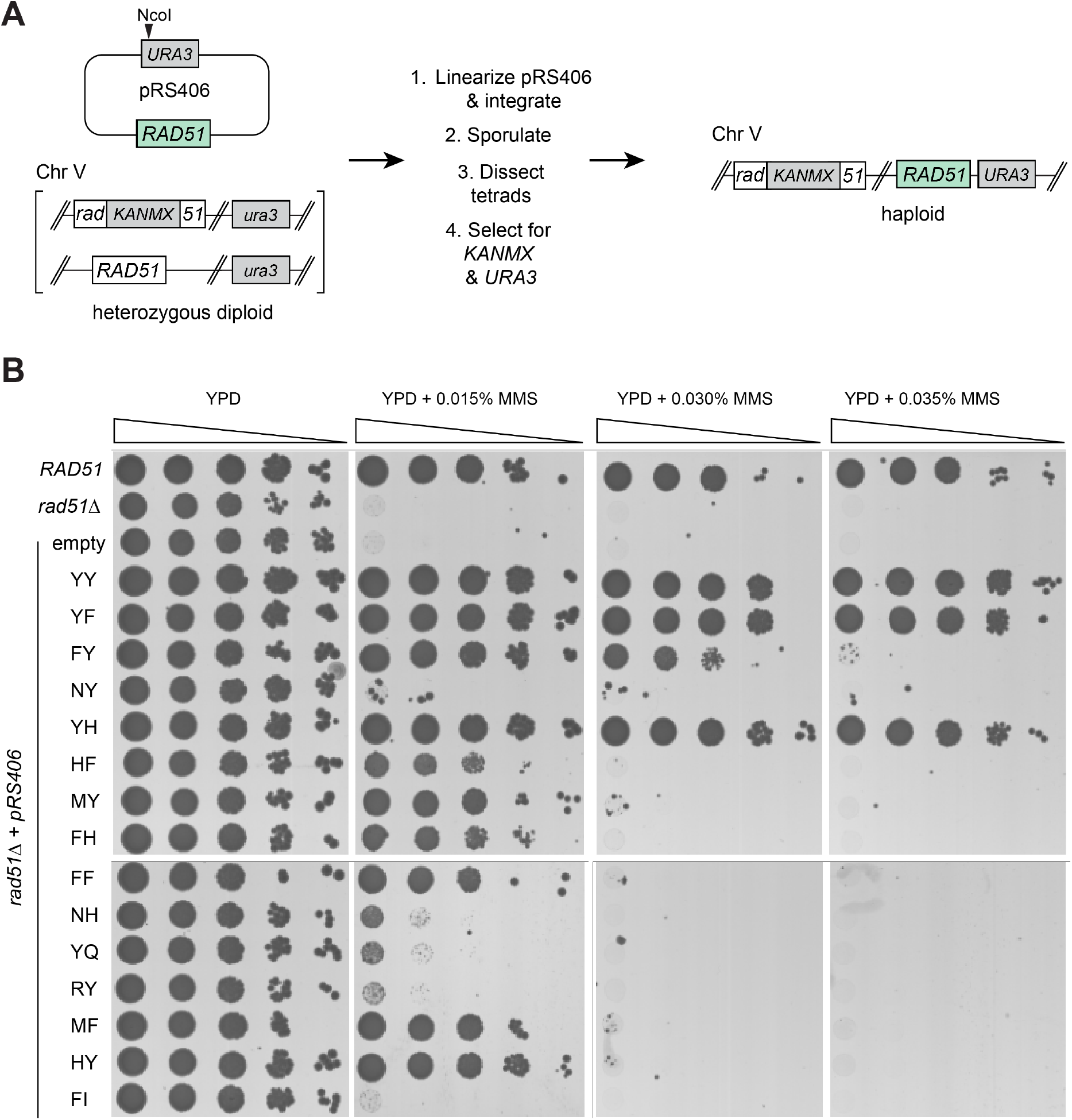
Genetic analysis of naturally occurring Rad51 variants. a, Schematic of method used to obtain haploid *S. cerevisiae* strains with *rad51* mutants integrated into chromosome V. **b,** Spot assays using 10-fold serial dilutions of each indicated strain. Cells were grown on YPD media with either no MMS, or with 0.015%, 0.03% or 0.035% MMS, as indicated. Wild-type *RAD51*, *rad51τι*, and empty vector controls are indicated.

Most of the naturally occurring alleles functioned within the context of *S. cerevisiae* Rad51 (11 out of 15, 73% of tested alleles) albeit to differing extents (Fig. 4b). Cells expressing three of the alleles grew well at the highest concentration of MMS (0.035%) used in these assays; these included the native *S. cerevisiae* allele YY, as expected, as well as the YF and YH alleles (Fig. 4b). YF was the second most abundant Rad51 allele identified in our bioinformatic analysis, corresponding to 35.7% of the analyzed sequences (including human RAD51), whereas YH was only found in 1.1% of the native Rad51 sequences (Fig. 3F). In addition, cells with the FY allele, which was the third most abundant allele in nature, corresponding to 11.5% of the native Rad1 sequences, grew at the next highest concentration of MMS (0.030%; Fig. 4b). Cells expressing six additional Rad51 alleles survived the lowest concentration of MMS tested (0.015%), these included HF, MY, FH, FF, MF, and HY (Fig. 4b). Lastly, cells expressing the NY, NH, YQ, RY alleles grew slightly better than the control *rad51τι* strain at 0.015% MMS, whereas cells with the FI allele closely resembled the *rad51τι* strain (Fig. 4b).

Taken together, our data show that many of the Rad51 sequence variants found in nature, corresponding to the same locations of the stacked tyrosines found in *S. cerevisiae* Rad51, still retain biological function with respect to MMS resistance when placed within the context of *S. cerevisiae* Rad51. The most robust alleles all correspond to amino acid residues capable of forming stacked pairs of aromatic side chains, and include YY, YF, FY and YH, whereas the second tier of functional alleles is expanded to include HF, MY, FH, FF, MF, and HY. The imidazole side chain of histidine commonly interacts with other aromatic residues (F, Y and W) via cation-ρε interactions, ρε-ρε stacking interactions, hydrogen-ρε interactions, or hydrogen bonding interactions (37, 38). Thus, with the exception of methionine (M), all of the functional alleles harbor aromatic residues capable of forming stacking interactions. The presence of methionine at the Rad51 interface can be explained by considering that methionine has a long unbranched side chain with varied functional groups that can adopt multiple configurations, thus offering the potential for ample structural plasticity, and methionine is well-known to interact with aromatic residues forming what is termed an S-aromatic contact (39, 40). Nevertheless, the limited sequence variation found in nature together with our experimental data using the natural sequence variants further reinforces the notion that the functional landscape of the Rad51 YY positions is highly restricted.

### The functional landscape for Rad51 interfacial amino acid residues

Next, we initiated a broader unbiased screen using deep mutagenesis to identify all possible sequence variations that might be tolerated at the *S. cerevisiae* Rad51 Y112 & Y253 positions. For this, we generated a library of *rad51* mutants encoded within a CEN plasmid (*pRS414–ScRAD51*) in which the codons for Y112 and Y253 were randomized to NNN, where N could be A, G, C, or T, yielding an input library with 400 hundred different pairwise amino acid combinations, plus mutants in which either codon was mutated to a stop codon. All functional *rad51* variants were identified based upon their ability to support growth of a *rad51Δ* strain on media containing 0.015% MMS (Fig. 5a). The resulting pool of survivors were analyzed by next generation DNA sequencing. Fold-enrichment for each variant was calculated by comparing the relative abundance of each allele recovered from cells grown in the absence of MMS versus cells grown the presence of MMS (Fig. 5b-5d). All experiments were done in three replicates, the mean fold-enrichment data are shown in Fig. 5, and data for each of the individual replicates are shown in Fig. S3 & Supplementary data file. Notably, the YY allele was highly enriched in the survivor pool (39-fold), providing an internal positive control demonstrating that the deep mutagenesis screen could recover the *S. cerevisiae* wild-type sequence (Fig. 5b- d). We did not recover alleles bearing stop codons at either position, providing an internal negative control confirming that full-length Rad51 is necessary for cell survival and growth on MMS (Fig. 5d). Moreover, there was good agreement between the alleles recovered in the deep mutagenic screen and the naturally occurring *RAD51* alleles that retained function when tested in *S. cerevisiae* (c.f. Fig. 4b & Fig. 5d); the sole exceptions were the HF and HY alleles which supported cell growth at 0.015% MMS (but not at higher MMS concentrations) when integrated into chromosome V, but were not enriched in the deep mutagenesis screen (c.f. Fig. 4b & Fig. 5d).

**Figure 5.**
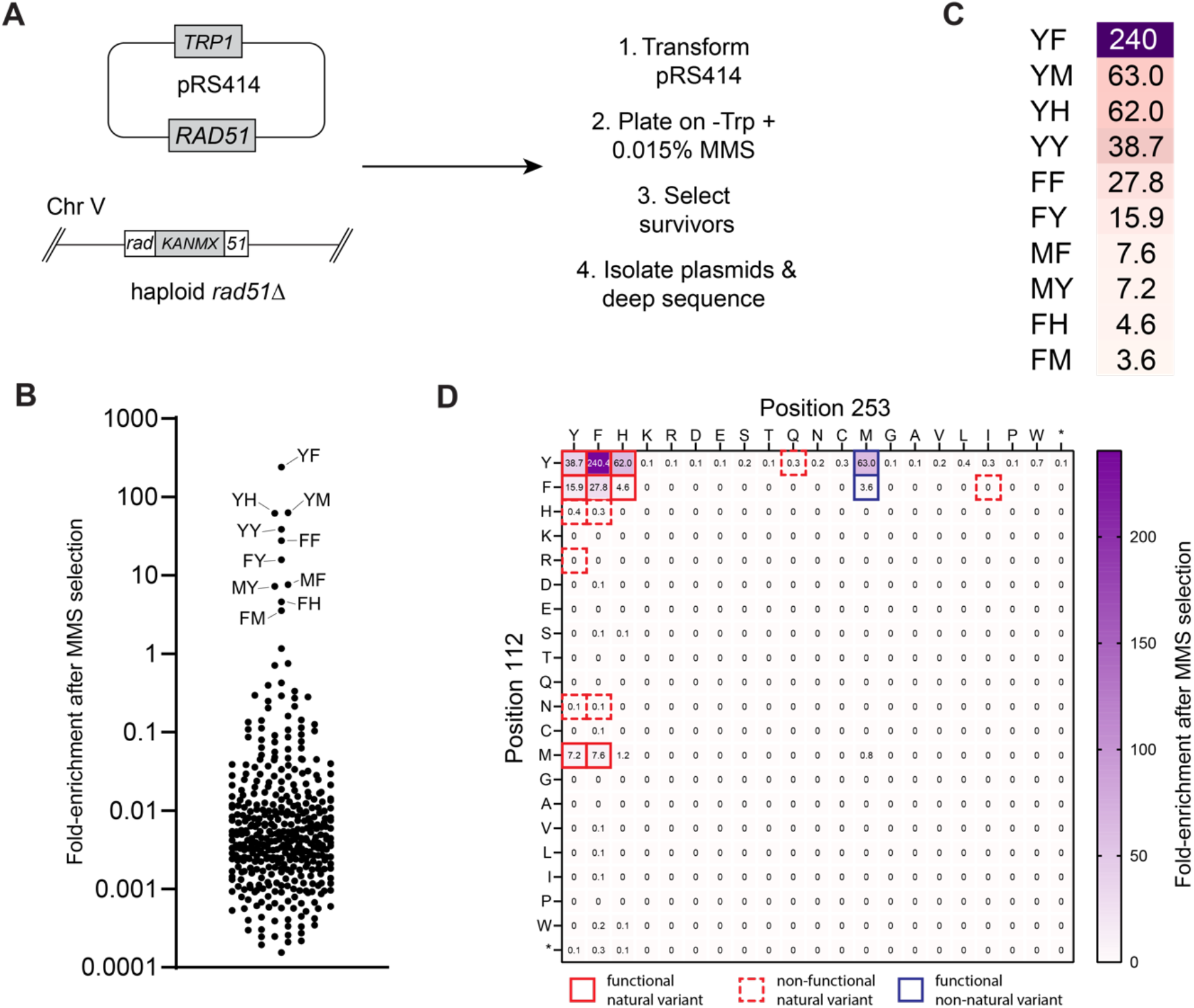
Deep mutagenic screen for all functional Rad51 interfacial amino acid residue pairs. a, Schematic of the high-throughput screen for *rad51* variants that allow for cell survival on media containing 0.015% MMS. **b,** Scatter plot showing the fold-enrichment of each *rad51* variant after selection with 0.015% MMS; the amino acid residue pairs of the ten most enriched *rad51* variants are highlighted; the data represent the mean values obtained from three separate screens (see Fig. S3 & Supplementary data file). **c,** Heat map legend highlighting the top ten most enriched *rad51* variants obtained in the deep mutagenic screen. **d,** Heat map showing the fold-enrichment for all pairwise interfacial amino acid residue combinations; stop codons are indicated with an asterisk; the data represent the mean values obtained from three separate screens (see Fig. S3 & Supplementary data file). Solid red boxes highlight functional natural sequence variants; dashed red boxes indicate non-functional natural variant sequences; blue boxes highlight functional non-native variant sequences.

Remarkably, of the 400 possible amino acid combinations, only 10 functional alleles were recovered with enhanced fold-enrichment after growth on MMS-containing media, corresponding to just 2.5% of the input library (Fig. 5b-d). Only four amino acid residues were represented within the functional Rad51 alleles (Y, F, H and M) whereas the remaining sixteen amino acid residues were all excluded from the pooled survivors (Fig. 5b-d). Moreover, the Rad51 deep mutagenesis screen did not recover an *S. cerevisiae* “Dmc1-like” SL allele, consistent with our other genetic assays. These findings further demonstrate that the functional landscape for amino acid residues at Rad51 positions 112 and 253 is highly restricted. Of the 15 natural variants tested above (Fig. 4), only 8 showed significantly enhanced enrichment in the deep mutagenesis screen (≥4-fold; YY, YF, YH, FY, FF, FH, MY, and MF), whereas the remaining 7 did not show any substantial enrichment (YQ, HY, HF, FI, RY, NY ad NH), suggesting that these alleles yielded nonfunctional proteins (Fig. 5b-d); it should again be noted that three of these nonfunctional natural alleles are exceedingly rare in nature, and may in fact reflect sequencing errors (YQ, 2 species; FI, 1 species, RY, 1 species). We speculate that the remaining natural variants not showing enhanced enrichment (*i.e.* HY, HF, NY, and NH) may originate from species with Rad51 proteins bearing compensatory mutations that allow these amino acids pairs to be more easily tolerated.

Interestingly, the YH allele was enriched to 62-fold, comparable to the native *S. cerevisiae* YY allele (39-fold enrichment; Fig. 5b-d), even though the YH allele is only found in 1.1% of the annotated *RAD51* genes (Fig. 3F). Similarly, the FF allele was enriched 28- fold (Fig. 5b-d), even though it is only found in 0.7% of the annotated *RAD51* genes (Fig. 3F). Likewise, the MY and MF alleles were enriched 7.2- and 7.6-fold (Fig. 5b-d), respectively, and were identified in 0.7% and 0.1% of the annotated *RAD51* genes (Fig. 3F). Whereas FY, the second most abundant native allele, was enriched by 16-fold (Fig. 5b-d). Surprisingly, the YF allele was enriched 240-fold (Fig. 5b-d); this allele is the second most commonly found in nature (Fig. 3F) and corresponds to the amino acid residue pair found in human RAD51. Two non-native alleles, YM (63-fold enrichment) and FM (3.6- fold enrichment), were also recovered in the survivor pool (Fig. 5b-d). Clearly these two alleles yield functional proteins capable of supporting cell growth in the presence of MMS, but their absence in nature suggests the possibility that there may be other functional constraints on Rad51 that are not fully revealed in the MMS assays.

These findings reinforce the notion that stacked aromatics (Y, F, H) are highly preferred at the Rad51 interface, which is consistent with our structural and bioinformatic analysis. Stacked phenylalanine (F) residues yield a functional protein even though the FF pair is only found in 0.7% of the native Rad51 sequences. In addition to the aromatic residues, methionine (M) also works well at the Rad51 interface, likely due to the conformational plasticity of the methionine side chain, its ability to interact with aromatic residues, or a combination of both properties. Although aromatic residues are highly preferred, there is a notable absence of tryptophan (W) in the native Rad51 sequences and tryptophan did not arise in the deep mutagenesis screen, even though from an evolutionary perspective tyrosine or phenylalanine mutations to tryptophan are considered conservative changes (41–43). We speculate that the tryptophan side chain may clash with surrounding residues in Rad51, or tryptophan may cause protein misfolding if exposed on the protein surface.

### Genetic analysis of natural Dmc1 sequence variants

The stacked aromatic interaction observed in the Rad51 lineage of the Rad51/RecA family of recombinases is absent from the Dmc1 lineage (Fig. 3). In the case of *S. cerevisiae* Dmc1 the corresponding residues are S48 and L189 (Fig. 3d). Bioinformatic analysis reveals that these residues in Dmc1 are most often MY (55.5%); GL (10.6%), SL (9.3%) or MF (6.2%), which together account for 81.6% of observed pairwise amino acid residue combinations; the remaining sequence variants are found less frequently (Fig. 3g-h).

Similar to our approach with Rad51, we sought to assess the functional constraints on the amino acid residues found at the Dmc1 protomer interfaces by first asking whether the naturally occurring sequences variants found in our bioinformatic analysis could retain function when placed within the context of *S. cerevisiae* Dmc1 (Fig. 6). For this, we constructed a total of 16 sequence variants, which together with the wild-type SL pair found in *S. cerevisiae*, account for the top 17 most common pairs of amino acids occurring at these positions in Dmc1 across eukaryotes (Fig. 3g). Genes encoding these alleles were cloned along with the promoter sequence of *DMC1* and integrated into chromosome V at the *URA3* locus in a strain deleted for endogenous *DMC1*. The resulting haploid yeast strains were mated to form diploid strains, and the diploid strains were assayed for meiotic progression, a phenotype that requires functional Dmc1; cells lacking Dmc1 exhibit cell cycle arrest in meiotic prophase I and nuclear divisions do not occur. Yeast strains harboring different *DMC1* alleles were made to synchronously enter meiosis, and meiotic progression was checked by scoring nuclear divisions at time points in sporulation conditions (Fig. 6a). As expected, the positive control with wild- type *DMC1* integrated into chromosome V exhibited meiotic progression similar to the WT strain, whereas the strains harboring an empty plasmid integration mimicked *dmc1τι* (Fig. 6b). Fourteen of the sixteen (87.5% of tested alleles) naturally occurring sequence variants that were tested within the context of *S. cerevisiae* Dmc1 retained biological function (Fig. 6b). This includes the wild-type *S. cerevisiae* SL pair and the Dmc1 variants containing MY, GL, MF, SV, AL, SQ, NH, GQ, AV, TA, NG, QF, and AQ in place of the wild-type SL amino acid residues (Fig. 6b). These findings further suggest that in contrast to Rad51, which shows a marked preference for stacked aromatic residues, the functional landscape of positions S48 and L189 in the Dmc1 monomer-monomer interface appears to be tolerant of a broader range of amino acid residues, although stacked aromatics are notably absent. Only two of the natural alleles, QW and MW, exhibited substantial meiotic defects (Fig. 6b). Both of these defective alleles contain tryptophan (W), and we speculate that the tryptophan side chain may either be too large to be easily accommodated within the specific context of *S. cerevisiae* Dmc1 or it may prevent proper protein folding.

**Figure 6.**
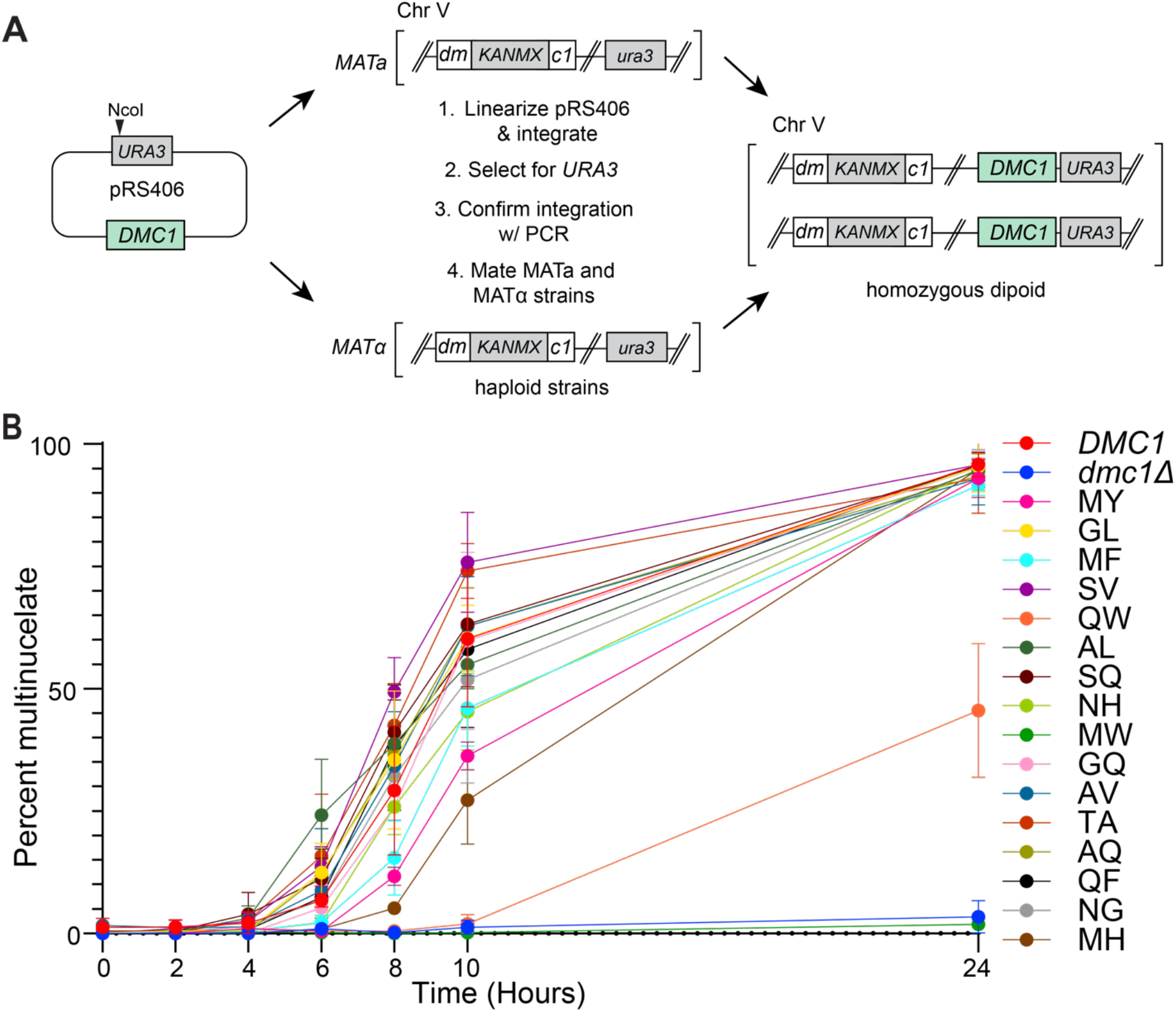
Genetic analysis of naturally occurring Dmc1 variants. a, Schematic of methodology used to obtain homozygous diploid *S. cerevisiae* strains with *dmc1* variants integrated into chromosome V. **b,** Graphical representation showing the percentage of multinucleate cells at each indicated time point after inducing sporulation. Each data point represents the mean and standard deviation of three separate assays.

### The functional landscape of Dmc1 interfacial amino acid residues

We next developed a deep mutagenesis screen to identify all possible sequence variations that might be tolerated at the *S. cerevisiae* Dmc1 S48 & L189 positions. For this, we generated a library of *dmc1* mutants, cloned along with the endogenous *DMC1* promoter sequence, encoded within a CEN plasmid (*pRS414–ScDMC1*) in which the codons for S48 and L189 were randomized. All functional *DMC1* variants were identified based upon their ability to support meiotic progression and undergo sporulation. Alleles that retained biological function were recovered by ether treatment, specifically selecting for ether-resistant *S. cerevisiae* spores while eliminating ether-sensitive vegetative cells that could not undergo sporulation (Fig. 7a)(44). The resulting pool of survivors were analyzed by next generation DNA sequencing. Fold-enrichment for each variant was calculated by comparing the relative abundance of each allele before and after ether selection (Fig. 7b-d). All experiments were done in three replicates, the mean fold- enrichment data are shown in Fig. 5, and data for each of the individual replicates are shown in Fig. S4 & Supplementary data file. The wild-type SL allele was recovered (2.2- fold enrichment), whereas stop codons were not, providing good internal positive and negative controls, respectively (Fig. 7b-d). In addition, 15 of the 17 natural *DMC1* variants that retained function when integrated into the chromosome (Fig. 6b) were also found to be functional in the deep mutagenesis assay (Fig. 7b-d), whereas the 2 alleles (QW & MW) that did not function when integrated into the chromosome (Fig. 6b) were not recovered in the deep mutagenesis screen. Of the 400 possible alleles, 182 appeared to retain at least some biological function (≥1-fold enrichment), corresponding to 45.5% of the input alleles (Fig. 7b-d). This result is in striking contrast to Rad51, where only 2.5% of the input alleles appeared to retain biological function. Remarkably, a total of 169 non-native alleles were recovered, highlighting the conformational plasticity of the Dmc1 interface (Fig. 7b-d). Surprisingly, the most enriched alleles were HM, TC, and NV, none of which are found in nature (Fig. 3h & Fig. 7-b). There were also examples of excluded amino acid residues, for example, at position 48 there was a striking tendency to avoid K, R, E, V, L, I, P, & M, whereas at position 189 there was a tendency to avoid the acidic residues D & E (Fig. 7d). In addition, although W was tolerated at position 48, this same residue was markedly absent from 189, with the sole exception of the WW pair (Fig. 7d). Lastly, there was some surprising overlap with the functional Rad51 variants, for example, Dmc1 alleles harboring YF & YH were both recovered in the screen (Fig. 7d). Moreover, as indicated above, we identified two functional Rad51 variants YM & FM, neither of which is found in nature, and both of these amino acid combinations were recovered in our screen for functional *DMC1* alleles (Fig. 7d). Taken together, these findings support the idea that the interfacial Dmc1 amino acid residues exhibit a high degree of functional and structural plasticity.

**Figure 7.**
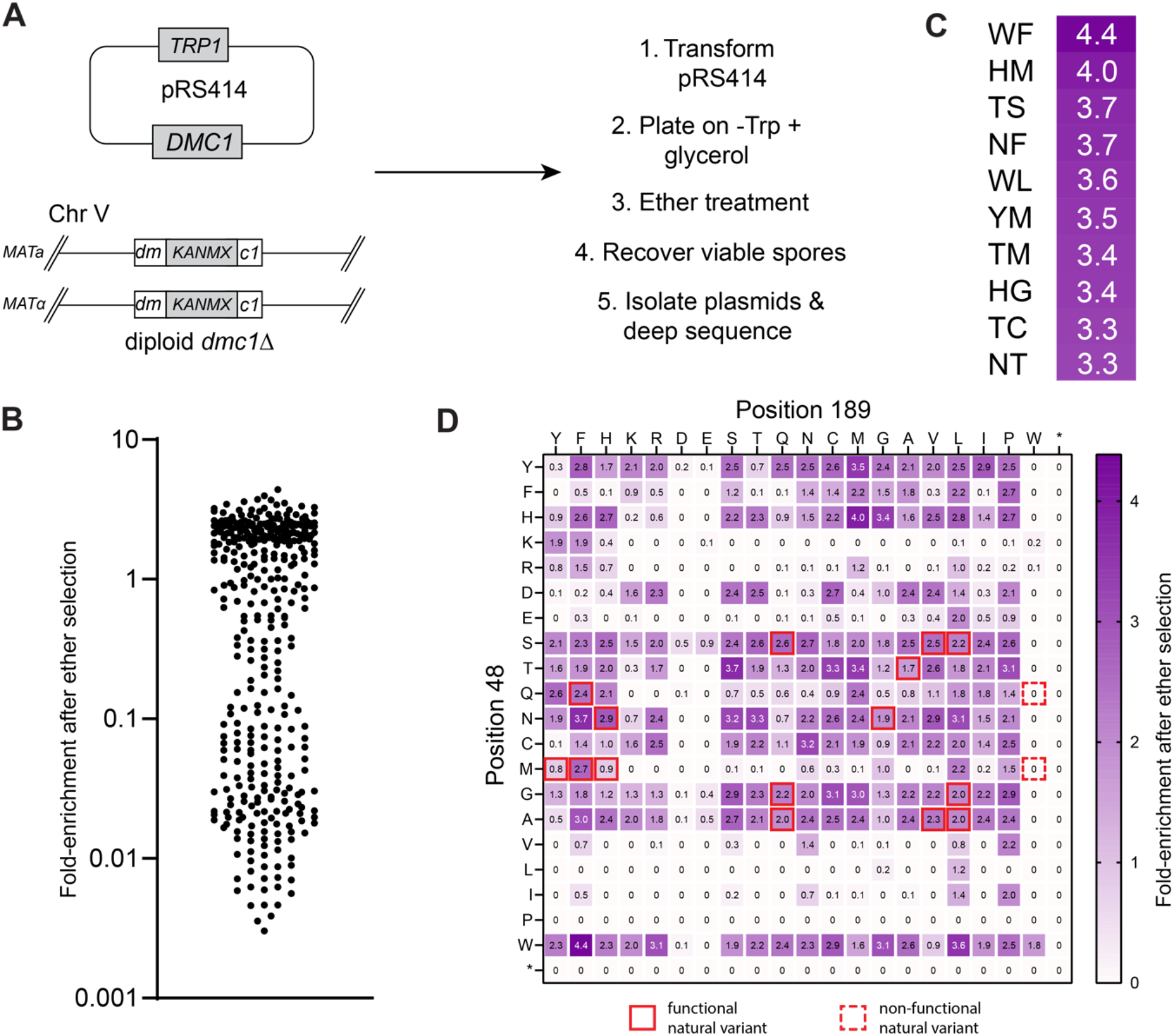
Deep mutagenic screen for all functional pairs of Dmc1 interfacial amino acid residues. a, Schematic of the high-throughput assay used to screen for functional *dmc1* variants. **b,** Scatter plot showing the fold-enrichment of all pairwise interfacial amino acid residue combinations; data reflect the mean of three separate screens (see Fig. S4 & Supplementary data file). **c,** Heat map legend highlighting the top ten most enriched *dmc1* variants obtained in the deep mutagenic screen. **d,** Heat map showing the fold- enrichment for all Dmc1 pairwise interfacial amino acid residue combinations; stop codons are indicated with an asterisk; heat map data reflect the mean of three separate screens (see Fig. S4 & Supplementary data file). Solid red boxes highlight function natural sequence variants and dashed red boxes highlight non-functional natural sequence variants.

## DISCUSSION

Rad51 and Dmc1 are important proteins that participate in the key DNA pairing reactions that take place during homologous recombination, but they each function in distinct biological situations. Rad51 is ubiquitously expressed and participates in both mitotic and meiotic recombination, whereas Dmc1 is uniquely expressed in meiosis and only functions in meiotic recombination. We still do not fully understand the unique functional attributes of each protein that allow for these distinct biological functions. Yet despite these differences in biological function, at a fundamental level both proteins fulfill the same biochemical activities, namely they assemble into helical filaments on ssDNA and promote stand invasion into a homologous DNA molecule.

Our data demonstrates that the stacked aromatic residues present at the interface between Rad51 monomers are crucial for biological function and reflect a highly conserved structural feature of the Rad51 lineage. In striking contrast, these stacked aromatic residues are absent from the Dmc1 lineage of the Rad51/RecA family of DNA recombinases. Although our experimental work utilized *S. cerevisiae* Rad51 and Dmc1, our findings are likely of relevance in other eukaryotes, given that these particular structural features are widely conserved. The functional significance of the YY residues found in *S. cerevisiae* Rad51 is highlighted by our finding that when they are mutated to the SL amnio acid residue pair found in Dmc1, the resulting “Dmc1-like” Rad51 protein is not functional *in vivo* and fails to assemble into nucleoprotein filaments on ssDNA *in vitro*. Moreover, our deep mutagenic scanning experiments show that the functional landscape of the YY pair positions is highly constrained, yielding only ten functional alleles most of which were stacked aromatics or aromatic residues in combination with methionine.

The presence of these highly conserved stacked aromatic residues within the protein-protein interfaces of the Rad51 filament raises the question how they might contribute to protein function. It should be noted that the presence of stacked tyrosine residues at the Rad51 interface was originally noted when the first crystal structure of *S. cerevisiae* was solved, and it was suggested that these residues might serve as sites of regulatory post-translational phosphorylation (34). However, we do not currently favor this hypothesis because there is no direct evidence that these residues are phosphorylated, there are no tyrosine-specific kinases in fungi (45), and amino acid residues from either native alleles or non-native alleles harboring F, H, or M would not be subject to the same post-translational mechanisms or modifications as Y, yet these alleles still retain biological function. Moreover, the observation that yeast Rad51-FF supports cell survival in the MMS assays would seem to suggest that if phosphorylation does take place, then it must not play a role in the Rad51-mediated repair of MMS-induced DNA damage.

Other potential roles for the stacked aromatic residues may be to modulate Rad51 filament stability, contribute to allosteric communication within the filaments, allow for self-recognition, or perhaps contribute to interactions with Rad51-specific filament end-binding factors. With respect to filament stability, our results with the “Dmc1-like” Rad51-SL mutant and the Rad51-EK salt bridge mutants provide clear evidence that disruption of the YY pair yields proteins that fail to assemble into stable filaments *in vitro*. Allosteric communication has been suggested as a means of coordinating ATP hydrolysis and strand exchange activities within the recombinase filaments albeit through poorly understood mechanisms (18, 34, 46–48), so it is formally possible that in the case of Rad51 this communication is regulated at least in part through the stacked aromatic residues. It is also possible that the stacked aromatic residues found in Rad51 may contribute to its ability to distinguish between self and non-self. We have previously shown that when mixed together, *S. cerevisiae* Rad51 and Dmc1 do not assemble into heterogenous intermixed filaments but rather assemble into homotypic side-by-side filaments on the same ssDNA molecules, illustrating that these proteins have an inherent propensity for self-recognition (49). This finding is also consistent with *in vivo* microscopy studies that have suggested the existence of side-by-side Rad51 and Dmc1 filaments in both *S. cerevisiae* and *Arabidopsis thaliana* (50–53). Lastly, it is possible that the stacked aromatic residues in Rad51 may contribute to interactions with Rad51- or Dmc1-specifc protein factors that may interact with the filament ends. In *S. cerevisiae*, potential examples of Rad51-specific interacting factors include, the antirecombinase Srs2 (54, 55), the recombination mediator protein Rad52 (23, 56–58), the Rad51 paralog complex Rad55- Rad57 (59), and the meiosis-specific Rad51 inhibitor protein Hed1 (60–63). Dmc1-specific factors include the meiosis specific protein complexes Hop2-Mnd1, which stimulates Dmc1-mediate strand exchange (62, 64–67), and the mediator complex Mei5-Sae3, which promotes Dmc1 filament assembly (14, 56, 68–70).

In addition to *Saccharomyces cerevisiae*, stacked aromatic residues are also found at the Rad51 interface for several important model organisms, including *Homo sapiens* (YF; Fig. S5A), *Mus musculus* (YF), *Gallus gallus* (HF), and *Schizosaccharomyces pombe* (YY). In striking contrast, Dmc1 lacks strongly conserved stacked aromatic residues at the monomer-monomer interface, which is instead most commonly occupied by MY, GL, or SL, which together account for approximately 75% of the *DMC1* genes that have been annotated. For example, as discussed above, in *S. cerevisiae* the YY residues found in Rad51 are replaced by SL in Dmc1. Similarly, stacked aromatics are also absent from Dmc1 in *Homo sapiens* (MY; Fig. S5B), *Mus musculus* (MY), *Gallus gallus* (MY), and *Schizosaccharomyces pombe* (MY).

Interestingly, although many organisms are thought to have both *RAD51* and *DMC1*, there are several examples of important model organisms that only have a *RAD51* gene and have lost the gene for *DMC1*. These include the ascomycetes *Sordaria macrospora* (YY), *Neurospora crassa* (YY), and *Podospora anserina* (YY), the pathogenic fungus *Ustilago maydis* (FY), and ecdysozoans such as *Caenorhabditis elegans* (FY; Fig. S5C) and *Drosophila melanogaster* (NH; Fig. S5D). In general, Rad51 from the twenty-eight *Drosophila sp.* available in our data set have mixed characteristics with 9 species harboring the more canonical Rad51 stacked aromatic residues (YY, YH, FH) and the remaining 19 species harboring alternative amino acid residues (NY or NH). Thus, with the exception of *Drosophila*, the model organisms that lack *DMC1* appear to have *RAD51* genes that follow the stacked aromatic pattern found in “canonical” Rad51 from organisms bearing both *RAD51* and *DMC1*. This exception may be related to the observation that *Drosophila* species are considered to have more rapidly evolving *RAD51* genes compared to other organisms (27).

The Rad51 and Dmc1 lineages of the Rad51/RecA family of recombinases are thought to have arisen from an ancient archaeal RadA progenitor. Interestingly, in archaeal RadA, the most common amino acids found at the position equivalent to the Rad51 stacked aromatics are VF (61%) and VY (15.1%); examples of prominent archaeal model organisms include *Haloferax volcanii* (VF), *Pyrococcus furiosus* (VF; Fig. S5), *Sulfolobus solfataricus* (VI), and *Methanococcus jannaschii* (TY). Thus, this pair of amino acid residues at the monomer-monomer interface of RadA are more similar to those found in Dmc1 in that there is not a strongly conserved pair of stacked aromatic residues. It is possible that the ancient RadA progenitor that gave rise to the Rad51 and Dmc1 lineages may have lacked the stacked aromatics, suggesting that stacked aromatic residues may be a more recent acquisition unique to the Rad51 lineage. Lastly, it should be noted that this interface in bacterial RecA is structurally distinct from that of the eukaryotic recombinases (Fig. S5F), so the protein-protein interfaces observed in the Rad51 and Dmc1 lineages appear to have arisen later in the evolutionary history of the Rad51/RecA family of recombinases.

## AUTHOR CONTRIBUTIONS

Y.S. conducted all work related to the CryoEM structures of Rad51 and Dmc1, and D-loop and single molecule assays. M.P. conducted all bioinformatic analysis, and designed and implemented the deep mutagenesis screens, and assisted with the genetic analysis of native Dmc1 variants. A.M.J. and R.M. conducted experiments related to the genetic analysis of the native Rad51 variants. K.M. conducted the genetic analysis of native Dmc1 variants. S.Y.K. assisted with the cloning and purification of Rad51 and Dmc1 proteins for CryoEM analysis and Rad51 mutants for biochemical assays. V.B.R provided guidance and assistance with all genetic assays. E.C.G. wrote the manuscript with assistance from all co-authors.

## ACKNOWLEDGMENTS

We thank Rodney Rothstein, Lorraine Symington, Andreas Hochwagen, and Luke Berchowitz for yeast strains and access to equipment. We thank Greene laboratory members for critically reading the manuscript. This research was funded by NIH grant R35GM118026 and NSF Grant MCB-1817315 (to E.C.G.). A.M.J., K.M., R.M., and S.K. were partially supported by the Columbia University Summer Undergraduate Research Fellowships (SURF) program and the Columbia College Summer Funding Program. This research was funded in part through the NIH/NCI Cancer Center Support Grant P30CA013696 and used the Genomics and High Throughput Screening Shared Resource. The CryoEM was in part supported by the National Cancer Institute’s National Cryo-EM Facility at the Frederick National Laboratory for Cancer Research under contract 75N91019D00024, some of the work was performed at the National Center for CryoEM Access and Training (NCCAT) and the Simons Electron Microscopy Center located at the New York Structural Biology Center, supported by the NIH Common Fund Transformative High Resolution Cryo-Electron Microscopy program (U24 GM129539,) and by grants from the Simons Foundation (SF349247) and NY State Assembly, and some of this work was performed at the Columbia University Cryo-Electron Microscopy Center.

## MATERIALS & METHODS

### Proteins

ScRad51 was overexpressed in *E. coli* BL21 (DE3) Rosetta2 cells transformed with plasmid encoding 6XHis-SUMO-ScRad51. Cells were grown in 2L LB media containing 100 μg/ml carbenicillin and 35 μg/ml chloramphenicol at 37°C until OD_600_ = 0.6, induced with 0.5 mM IPTG and grown for 3 hours at 37°C. Cell paste was suspended in 50 ml lysis buffer (50 mM Tris-HCl [pH 7.5], 10% glycerol, 1M NaCl, 1mM DTT, 0.5 mM PMSF, 15 mM imidazole, 0.1% Tween 80, 1 protease inhibitor tablet) and lysed by sonication. The lysate was centrifuged at 35,000 rpm for 45 mins followed by precipitating the supernatant with 12g ammonium sulfate for 1 hour. The precipitate was spun down at 10,000 rpm for 30 mins. The pellet was dissolved in 50 ml binding buffer (25 mM Tris-HCl [pH 7.5], 10% glycerol, 200 mM NaCl, 0.1% Triton X-100, 15 mM imidazole, 5 mM beta- mercaptoethanol) and applied to the 5 ml HisPur^TM^ Ni-NTA resin (Thermo Fisher Scientific) equilibrated with the same binding buffer. The protein was eluted with 10 ml elution buffer (25 mM Tris-HCl [pH 7.5], 10% glycerol, 200 mM NaCl, 200 mM imidazole, 0.1% Triton X-100). SUMO protease was added to the elution, followed by dialyzing for 16 hours at 4°C in dialysis buffer (50 mM Tris-HCl [pH 7.5], 200 mM NaCl, 10% glycerol, 15 mM imidazole, 1 mM DTT). The sample was re-applied to the 5 ml Ni-NTA resin equilibrated with the binding buffer and flow-through was collected and concentrated to 30 μM. The protein was flash-frozen in liquid nitrogen and stored at –80°C

ScDmc1 was overexpressed in *E. coli* BL21 (DE3) Rosetta2 cells transformed with plasmid encoding 6XHis-ScDmc1. Cells were grown in 2L LB media containing 100μg/ml carbenicillin and 35 μg/ml chloramphenicol at 37°C until OD_600_ = 0.8, induced with 0.1 mM IPTG and grown for 16 hours at 16°C. Cell paste was suspended in 100 ml lysis buffer (50 mM Tris-HCl [pH 7.5], 10% glycerol, 500 mM KCl, 0.01% Triton X-100, 1mM DTT, 2 mM ATP, 2 mM MgCl_2_, 1mM PMSF, 1 protease inhibitor tablet) and lysed by sonication. The lysate was centrifuged at 35,000 rpm for 45 mins and the supernatant was applied to 5 ml Talon resin (Takara) equilibrated with the binding buffer (25 mM Tris-HCl [pH7.5], 10% glycerol, 150 mM KCl, 0.01% Triton X-100, 2 mM ATP, 2 mM MgCl_2_). The resin was washed with wash buffer (25 mM Tris-HCl [pH7.5], 10% glycerol, 500 mM KCl, 0.01% Triton X-100, 2 mM ATP, 2 mM MgCl_2_) followed by re-equilibrating with the binding buffer. The protein was eluted with 10 ml elution buffer (25 mM Tris- HCl [pH 7.5], 10% glycerol, 150 mM KCl, 200 mM imidazole, 0.01% Triton X-100, 2 mM ATP, 2 mM MgCl_2_) followed by dialyzing for 16 hours at 4°C in dialysis buffer (25 mM Tris-HCl [pH 7.5], 100 mM KCl, 10% glycerol, 0.01% Triton X-100, 2 mM ATP, 2 mM MgCl_2_, 0.5 mM EDTA). Next day, the sample was injected to a 1 ml heparin sepharose column (GE Healthcare) and fractionated with 100 – 600 mM KCl gradient. The fractions containing ScDmc1 were combined, concentrated to 80 μM, flash-frozen in liquid nitrogen and stored at –80°C.

**CryoEM sample preparation**

To prepare the *S. cerevisiae* Rad51 filaments for CryoEM analysis, purified ScRad51 (5 μM) was mixed with 0.25 µM of a 96–mer ssDNA (IDT; 5’– AAT TCT CAT TTT ACT TAC CGG ACG CTA TTA GCA GTG AAA ATT TCC TGA TAG TCG TCA CCG CGT TTT GCG CAC TCT TTC TCG TAG GTA CTC AGT CCG–3’) in HR buffer (30 mM HEPES [pH 7.5], 50 mM KCl, 20 mM MgCl_2_, 1 mM DTT and 2 mM ATP) and incubated at 30°C for 15 minutes. A sample volume of 3.5 µl was applied to a glow-discharged UltrAuFoil grid (R 0.6/1, 300 mesh), blotted for 4 seconds and plunge-frozen in liquid ethane using Vitrobot Mark IV (FEI, USA) at 100% humidity and 4°C. To prepare the *S. cerevisiae* Dmc1 filaments for CryoEM analysis, purified ScDmc1 (5 μM) was mixed with 0.25 µM of 96– mer ssDNA (as above) in HR buffer supplemented with 1.5 mM CaCl_2_ and the sample was incubated at 30°C for 5 minutes. The samples were then supplemented with 8 mM CHAPSO (3-([3-cholamidopropyl]dimethyammonio)-2-hydroxy-1-propanesulfonate; Hampton Research) prior to being applied to grids and plunge-frozen in liquid ethane, as described above for Rad51.

### CryoEM data acquisition

Samples were initially screened using a Glacios (Thermo Fisher, 200 keV). Grids selected for high-resolution data collection were imaged using a Titan Krios (Thermo Fisher) microscope equipped with a K3 direct electron detector (Gatan). Grids were imaged at 300 keV in electron counting mode, with a defocus range of -1.0 μm to -2.5 μm. 0.846 Å and 0.83 Å image pixel sizes and a nominal dose of 51.19 e^-^/Å^2^ and 58 e^-^/Å^2^ were used for Rad51 and Dmc1, respectively.

### CryoEM data processing

Data was processed using cryoSPARC v4.3.1 (71). The movies were aligned and dose-weighted using Patch-motion correction and Patch-CTF estimation (72, 73). Particles were picked, extracted, and classified to exclude junk particles and generate maps of Rad51-DNA filament or Dmc1-DNA filament ab initio. The class corresponding to the Rad51 filament or Dmc1 filament were selected and refined by optimal per-particle defocus using local CTF refinements. The nominal resolution of the CryoEM map was estimated by 0.143 gold standard Fourier Shell Correlation (FSC) cut off. To refine the Rad51 and Dmc1 filament structures, the human RAD51 structure (PDB: 5H1B)(36) and the predicted yeast Dmc1 structure from the AlphaFold database (74, 75), respectively, were fit into the corresponding CryoEM density maps using Chimera (76), followed by manually correcting and fitting amino acid residues to the density map using Coot (77). The structure was real-space refined by using rigid-body refinement, secondary structure, Ramachandran, rotamer and reference model restraints in Phenix (78).

### D–loop assays

Presynaptic complexes were assembled by incubating Rad51 (300 nM) (wild-type or mutants) and 90-mer ssDNA (10 nM) at 30°C for 15 min in the standard HR buffer (30 mM Tris-OAc, pH 7.5, 20 mM MgOAc, 50 mM KCl, 1 mM DTT, 0.2 mg/ml BSA) in the presence of 10 mM ATP. The resulting PSCs were then mixed with Rad54 (95 nM), RPA (750 nM), and plasmid that contains homology to the ssDNA (9.25 nM) and further incubated for 5 min to allow D-loop formation. The reactions were quenched by adding stop buffer and proteinase K and resolved on a 0.9% agarose gel in 1X TAE. D-loop products were imaged on GE Healthcare Life Sciences Typhoon FLA 9500 biomolecular imaging system.

### Single molecule assays

Experiments were conducted and analyzed as described previously (79). All assays were conducted at 30°C. Briefly, the ssDNA was tethered to the bilayer and aligned by flowing GFP-RPA (100 nM) with BSA buffer (40 mM Tris-HCl [pH8.0], 2 mM MgCl_2_, 1 mM DTT, 0.2 mg/mL BSA). Rad51 filament formation was initiated by injecting Rad51 (1 μM) in the HR buffer (30 mM Tris-Ac [pH 7.5], 50 mM KCl, 5 mM MgAc, 1 mM DTT, 0.3 mg/mL BSA, and 2 mM ATP). Rad51 filament assembly rate was measured based upon the loss of GFP-RPA signal.

### Genetic assays of Rad51 sequence variants

Wild type *RAD51*, along with 500 base pair upstream and downstream of the gene, was amplified and cloned into the yeast integrative vector pRS406 using PCR-based methods. *RAD51* point mutations were obtained by site directed mutagenesis. Vectors containing the *RAD51* gene (both wildtype and the mutants) were linearized at the *URA3* locus using NcoI-HF restriction digestion enzyme and transformed into a diploid yeast strain (*yECG48*) heterozygous for *RAD51 (MATalpha rad51Δ::KanMX; ADE2 leu2-3,112 his3- 11,15 ura3-1 lys2Δ / MATa TRP1 his3-11,15 leu2-3,112 ura3-1 lys2Δ ADE2 RAD5)*. Positive and single copy integrations into the host chromosome were confirmed by PCR.

Positively transformed yeast strains were then subjected to sporulation and tetrads dissected to obtain a haploid yeast strain harboring a *RAD51* allele at the *ura3* locus and deletion of *rad51* at the endogenous locus. Haploid strains having only one copy of *RAD51* in the genome were employed for the spot assays. For overexpression assays, haploid *RAD51* strains harboring the E, K, S and L mutations (as indicated) were transformed with point mutations of *RAD51* cloned into a 2 micron plasmid for and employed for spot assays. Spot assays were performed as follows: Yeast strains with indicated genotypes were grown in liquid YPD (or synthetic drop-out media, as indicated) overnight at 30°C. Cultures were diluted to OD_600_ of 1.0 in water the following morning. Serially diluted cultures (4 µL) were spotted on both freshly poured YPD (or synthetic drop-out plates) and YPD (or synthetic drop-out) plates containing the indicated amounts of methyl methanesulfonate (MMS; Sigma-Aldrich, Cat. No. 129925). The plates were incubated at 30°C and imaged 2-3 days post spotting.

### Deep mutagenesis screen for Rad51

The pool of *ScRAD51* genes with randomized codons corresponding to amino acid residues 112 and 253 in a centromeric plasmid (pRS414) was made by PCR amplification of *pRS414–ScRAD51* in two fragments using the following two sets of degenerate primers: (set 1) MTP7: 5’ – ACT GCT GAA GCG GTA GCA NNN GCT CCC AGA AAG GAT TTA TTG GAA – 3’ plus MTP8: 5’ – TAA CTG ATG ATC GGC GTT NNN GGC TCT TGC ATA CGC AAC G – 3’); and (set 2) MTP3: 5’ – AAC GCC GAT CAT CAG TTA AGA CTT – 3’ plus MTP4: 5’ – TGC TAC CGC TTC AGC AGT G – 3’. Following the PCR reactions, the samples were digested with DpnI (20 units; NEB, Cat. No. R0176), assembled using In-Fusion Snap Assembly (Takara Bio, Cat. No. 638947) and transformed into Stellar competent *E. coli* cells (Takara Bio, Cat. No. 636763). The transformed cells were plated onto LB plus 100 µg/ml carbenicillin and grown overnight at 37°C. The resulting transformants (100,000+ clones total) were collected in LB supplemented with 100 μg/mL carbenicillin, diluted 1:20 in the same media (to OD_600_ = 2-3), and incubated at 37°C with shaking for three hours before plasmids were extracted.

The degenerate pool was transformed into strain *yECG81* (*MATa rad51::URA3 lys2 leu2-3,112 trp1-1 ura3 his3-11,15 RAD5 ADE2*) using the lithium acetate method onto synthetic drop-out media lacking uracil (SD –Ura) as previously described (80). After a 72-hour incubation at 30°C, ≥10,000 transformants were collected, resuspended to a calculated OD_600_ = 10, and 100 µL were plated on SD –Ura supplemented with 0.015% MMS. After a 72-hour incubation at 30°C, ∼100,000 colonies were collected. Plasmids were extracted from cell populations before and after MMS selection using Wizard Plus SV Miniprep DNA Purification System (Promega), with the protocol modified as following: after addition of cell lysis buffer, 200 µL glass beads were added and the mixture was vortexed for a total of two minutes. The resulting DNA (20 ng) was used as template in a 50 µL PCR1 reaction using MTP37 (5’ – CTC TTT CCC TAC ACG ACG CTC TTC CGA TCT TGG GCT TCA CAC TGC TGA AG – 3’) plus MTP38 (5’ – CTG GAG TTC AGA CGT GTG CTC TTC CGA TCT CCA GAA GTC TTA ACT GAT GAT CGG C – 3’) with PrimeSTAR Max DNA Polymerase (Takara Bio) to amplify a portion of Rad51 with universal Illumina adaptor 5’ overhangs. Product from PCR1 was purified using Nucleospin Gel and PCR Clean-up (Takara Bio), and 10 ng was used as template for PCR2 reaction with indexed p5/p7 primers. Thermocycler conditions were as follows for PCR1: 98°C for 30 seconds, 98°C for 10 seconds, 60°C for 15 seconds, 72°C for 15 seconds (steps 2-4 repeated 25 times), 72°C for two minutes. For PCR2, the annealing temperature was 65°C and steps 2-4 were repeated 10 times. Barcoded PCR2 reactions were pooled, resolved by 1% agarose gel electrophoresis, and DNA was isolated by Gel Extraction Kit (Qiagen). Paired-end sequencing was performed using the NextSeq platform with automated demultiplexing and adaptor trimming (Illumina). Resulting reads contained paired 76 bp Rad51 sequence covering the 112 and 253 positions.

Reads were processed using BBTools (81) to fuse correct sized paired reads and remove reads lacking 20 bp sequence upstream and downstream both of the 112 and 253 position codons. 112 and 253 position codon sequences were extracted, translated, and counted using custom Python code. A fold-enrichment score was calculated by comparing the normalized frequency of each amino acid pair before and after MMS selection. Data was plotted as heatmaps using matplotlib/seaborn (82, 83).

### Genetic assays for natural Dmc1 variants

*DMC1* natural variant mutants were generated by site-directed mutagenesis with PCR amplification of *pRS406-ScDMC1* in two fragments with one of MTP21 through MTP26 (5’ –ACA CAG TCA ATA CCG TTT TGN NNA CAA CAA GAA GAC ATC TAT GTA AAA TTA AA–3’) plus MTP28 through MTP35 (5’ – TCC ATT TGA TGT TCA CTA TTN NNG GCT CTA GCA TAT GAA ACG TTT GC– 3’; NNN represents the location of the nucleotides encoding the randomized amino acid residue) and MTP13 (5’– AAT AGT GAA CAT CAA ATG GAA CTT GTT G–3’) plus MTP14 (5’– CAA AAC GGT ATT GAC TGT GTA TAT CCC–3’), followed by DpnI (NEB) digestion, In-Fusion Snap Assembly (Takara Bio), and transformation into Stellar competent cells (Takara Bio). Individual transformants were streak purified, plasmids extracted, and mutations confirmed by whole plasmid sequencing (Plasmidsaurus).

Plasmid DNA (1.3 μg) was digested with NcoI to linearize at the *URA3* gene and transformed into yECG008/AH11524 (SK1 *MATa ho::LYS2 ura3 leu2::hisG his3::hisG trp1::hisG dmc1::KanMX*) and yECG009/AH11525 (SK1 *MATalpha ho::LYS2 ura3 leu2::hisG his3::hisG trp1::hisG dmc1::KanMX*; the SK1 strains were a gift from Andreas Hochwagen) via the lithium acetate/ssDNA carrier method. Single copy integrations at the chromosomal *URA3* locus were confirmed by PCR. Haploid strain pairs for each natural variant were mated, diploids selected by colony morphology and confirmed by replica plate mating with SK1 *MATa* and *MATalpha* tester strains (tester strains were a gift from Luke Berchowitz).

Synchronous meiotic cultures were prepared as previously described (84). Diploid strains were patched from –80°C stocks onto YPG (1% yeast extract, 2% peptone, 2% glycerol) solid media and grown for 18 hours at 30°C. Resulting cell patches were re-patched onto YPD 4% glucose and grown for 24 hours at 30°C. Resulting patches were inoculated into YPD liquid cultures and grown overnight. YPD cultures were then diluted to a calculated OD_600_ = 0.3 in BYTA (1% yeast extract, 2% tryptone, 1% potassium acetate, 50 mM potassium phthalate) and grown for 20 hours. BYTA cultures were harvested, washed once in sporulation media (SPO, potassium acetate, 0.02% raffinose), and resuspended in SPO to a calculated OD_600_ = 1.8. Sporulation cultures were incubated at 30°C for 24 hours, with a portion of the culture being removed and fixed in 3.7% formaldehyde at 4°C overnight at time points indicated in the figures. Cells were resuspended in KPO_4_/sorbitol (1.2 M sorbitol, 100 mM potassium phosphate [pH 7.5]) plus 1% Triton-X and incubated for 5 minutes at room temperature, washed once in KPO_4_/sorbitol and resuspended in DAPI mounting solution (VECTASHIELD Antifade Mounting Medium with DAPI, Cat. No. H-1200). Cells were imaged on a Nikon Eclipse TE2000-U and the number of DAPI-stained nuclei were used to score a minimum of 200 cells per time point as mononucleate, dinucleate, or tetrads.

### Deep mutagenesis screen for Dmc1

The pool of *ScDMC1* genes with randomized amino acid residues 48 and 189 position codons in a centromeric plasmid was made by PCR amplification of *pRS414–ScDMC1* in two fragments using the following two sets of degenerate primers: (set 1) MTP17: 5’ – ACA CAG TCA ATA CCG TTT TGN NNA CAA CAA GAA GAC ATC TAT GTA AAA TTA AA – 3’ plus MTP18: 5’ – TCC ATT TGA TGT TCA CTA TTN NNG GCT CTA GCA TAT GAA ACG TTT GC– 3’); and (set 2) MTP13: 5’ – AAT AGT GAA CAT CAA ATG GAA CTT GTT G– 3’ plus MTP14: 5’ – CAA AAC GGT ATT GAC TGT GTA TAT CCC –3’. Following the PCR reactions, the samples were digested with DpnI (20 units; NEB, Cat. No. R0176), assembled using In-Fusion Snap Assembly (Takara Bio, Cat. No. 638947) and transformed into Stellar competent *E. coli* cells (Takara Bio, Cat. No. 636763). The transformed cells were plated onto LB plus 100 µg/ml carbenicillin and grown overnight at 37°C. The resulting transformants (100,000+ clones total) were collected in LB supplemented with 100 μg/mL carbenicillin, diluted 1:20 in the same media (to OD 2-3), and incubated at 37°C with shaking for three hours before plasmids were extracted.

The degenerate pool was transformed into strain yECG041 (SK1 *MATa ho::LYS2 ura3 leu2::hisG his3::hisG trp1::hisG dmc1::KanMX // SK1 MATalpha ho::LYS2 ura3 leu2::hisG his3::hisG trp1::hisG dmc1::KanMX*) or yECG010/A4962 (SK1 *MATa ho::LYS2 lys2 ura3 leu2::hisG his3::hisG trp1::hisG // SK1 MATalpha ho::LYS2 lys2 ura3 leu2::hisG his3::hisG trp1::hisG*; both strains were a gift from Andreas Hochwagen) using the lithium acetate method onto synthetic drop-out media lacking tryptophan or histidine (SD –Trp or SD – His) containing 2% glucose as previously described (80). After 72-hour incubation at 30°C, ≥10,000 transformants were collected in SD –Trp/–His plus 2% glucose, centrifuged at 3,000 x g for 1 minute, resuspended to a calculated OD_600_ = 200, and 150 µL were plated on SD –Trp/–His containing 2% glycerol to induce sporulation. After 72-hour incubation at 30°C, colonies were collected in sterile water and the cell solutions were transferred to a glass flask. An equal volume of diethyl ether (5 mL) was added, and the mixture was incubated at room temperature with gentle shaking for 15 minutes. After incubation, a sample of the cell-containing aqueous layer was removed, centrifuged at 20,000 x g for 1 minute, and resuspended in the same volume water. Samples of 100 µL were plated on SD –Trp plus 2% glucose to recover ether treatment survivors. After 72-hour incubation at 30°C, colonies were scaped into water and a portion of cells used for plasmid extraction. Plasmids were extracted before sporulation and either treatment (directly from transformation selection plate) and after ether treatment using Zymoprep Yeast Plasmid Miniprep II kit (Zymo Research, Cat. No. D2004).

---

**Figure S1.**
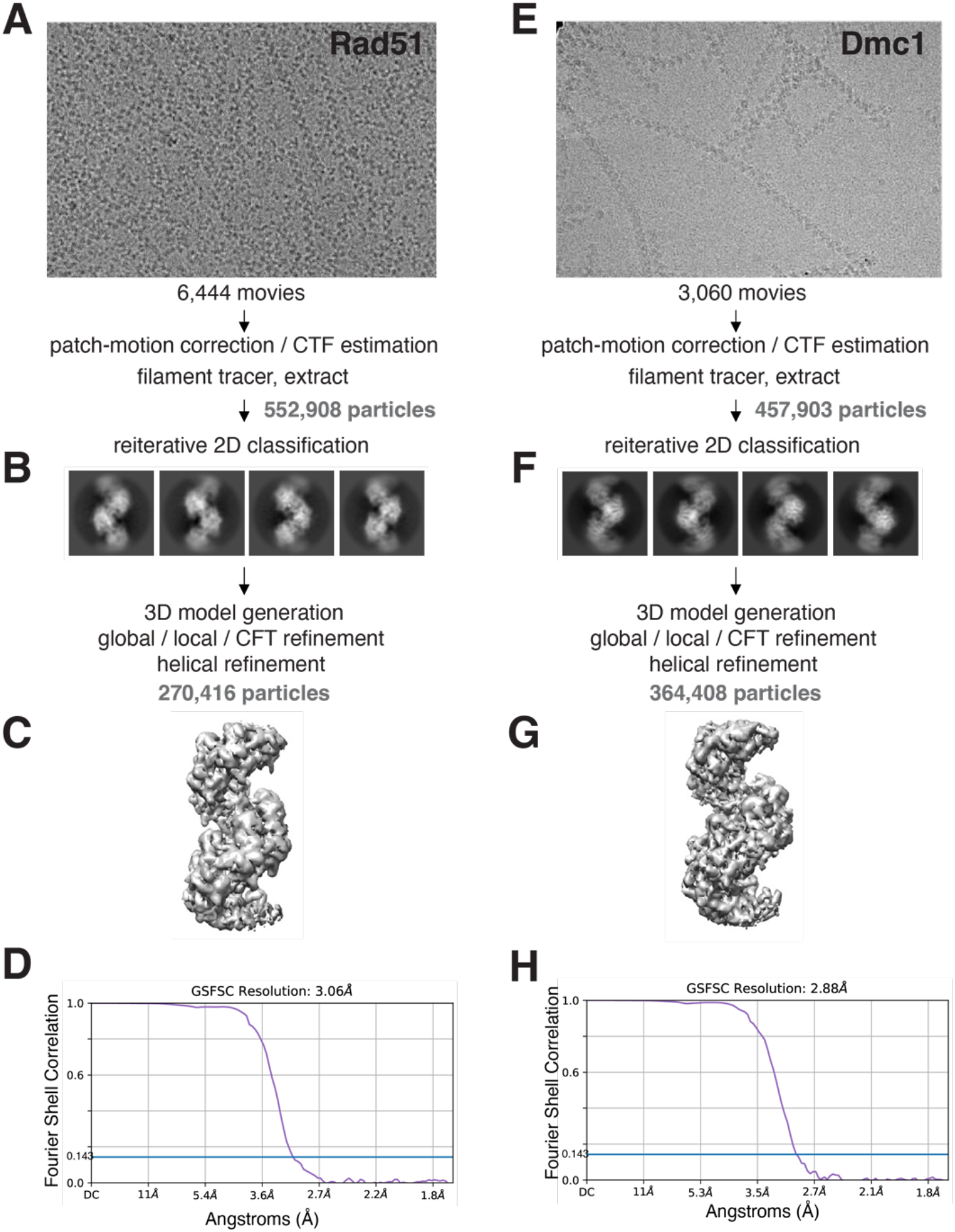
Cryo-EM data processing for the Rad51 and Dmc1 nucleoprotein filaments. a, Representative micrograph used for Rad51 filament data processing. **b,** Representative 2D classes. **c,** Final 3D map for the Rad51 filament. **d,** Fourier shell correlation curve for the Rad51 data. **e,** Representative micrograph used for Dmc1 filament data processing. **f,** Representative 2D classes. **g,** Final 3D map for the Dmc1 filament. **h,** Fourier shell correlation curve for the Dmc1 data.

**Figure S2.**
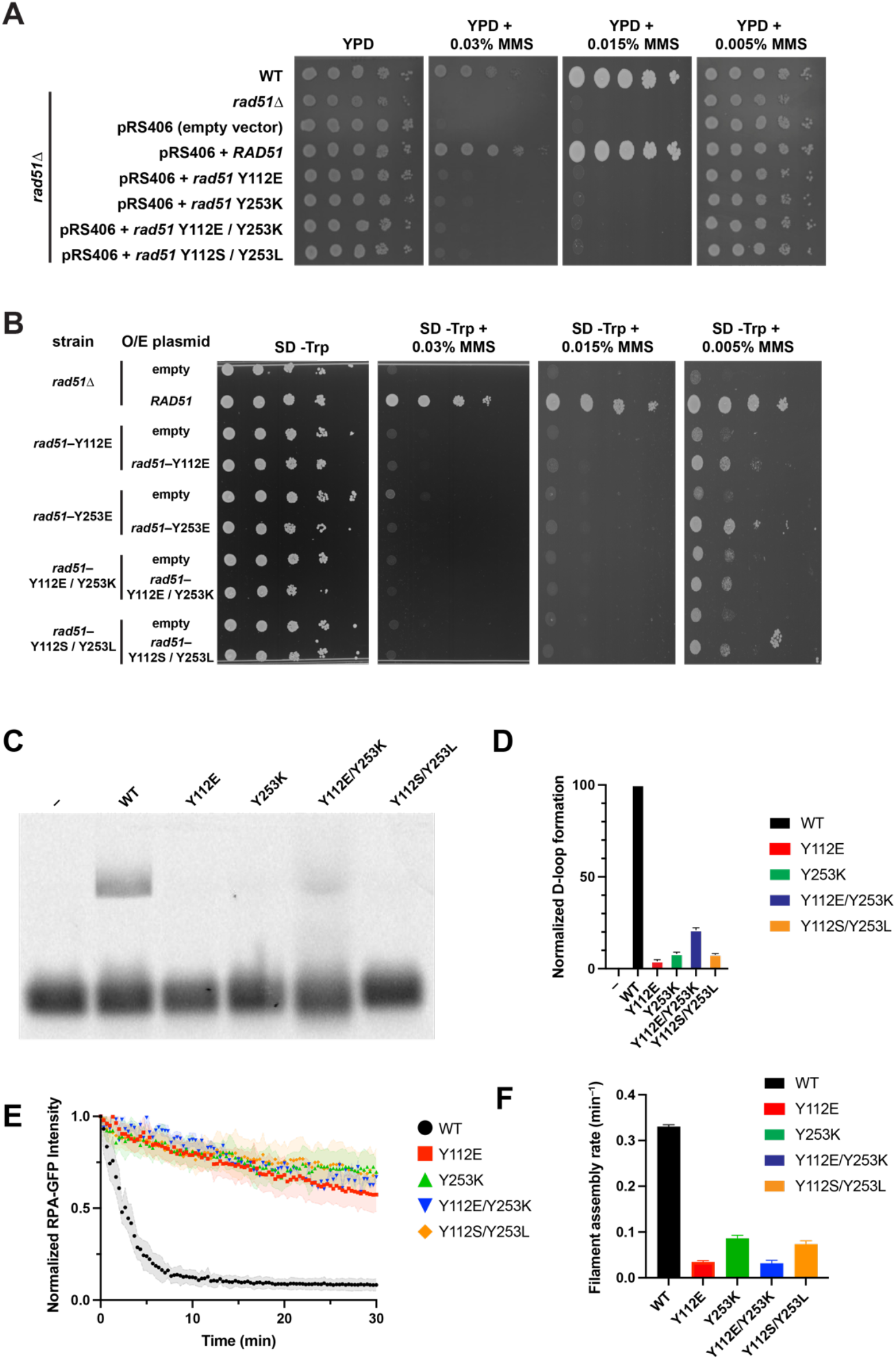
Genetic & biochemical analysis of YY to EK mutants. a, Spot assays using 10-fold serial dilutions of *S. cerevisiae* strains expressing *rad51* Y112E, Y253K, Y112E + Y253K or Y112S + Y253L, as indicated, and grown on YPD plus 0.03, 0.015, or 0.005% MMS. In each case, the indicated *rad51* mutant was integrated into chromosome V as depicted in Fig. 4a. **b,** Spot assays using the indicted *rad51* mutant strains plus a plasmid containing the same *rad51* gene to allow for protein overexpression (O/E) and grown on synthetic dropout media minus Tryptophan (SD -Trp) plus 0.03, 0.015, or 0.005% MMS, as indicated. **c,** D-loop assays with the indicated Rad51 proteins. **d,** Graphical representation of normalized D-loop formation. Data were all normalized to reactions with wild-type Rad51 and bars for the mutant proteins correspond to the mean and stand deviation of three separate experiments. **e,** Single molecule Rad51 filament assembly assays on ssDNA bound by GFP-RPA; the loss of GFP-RPA signal intensity is a proxy for Rad51 filament assembly. **f,** Quantitation of Rad51 filament assembly rates; bars correspond to the mean and stand deviation for n = 30 molecules for each experiment.

**Figure S3.**
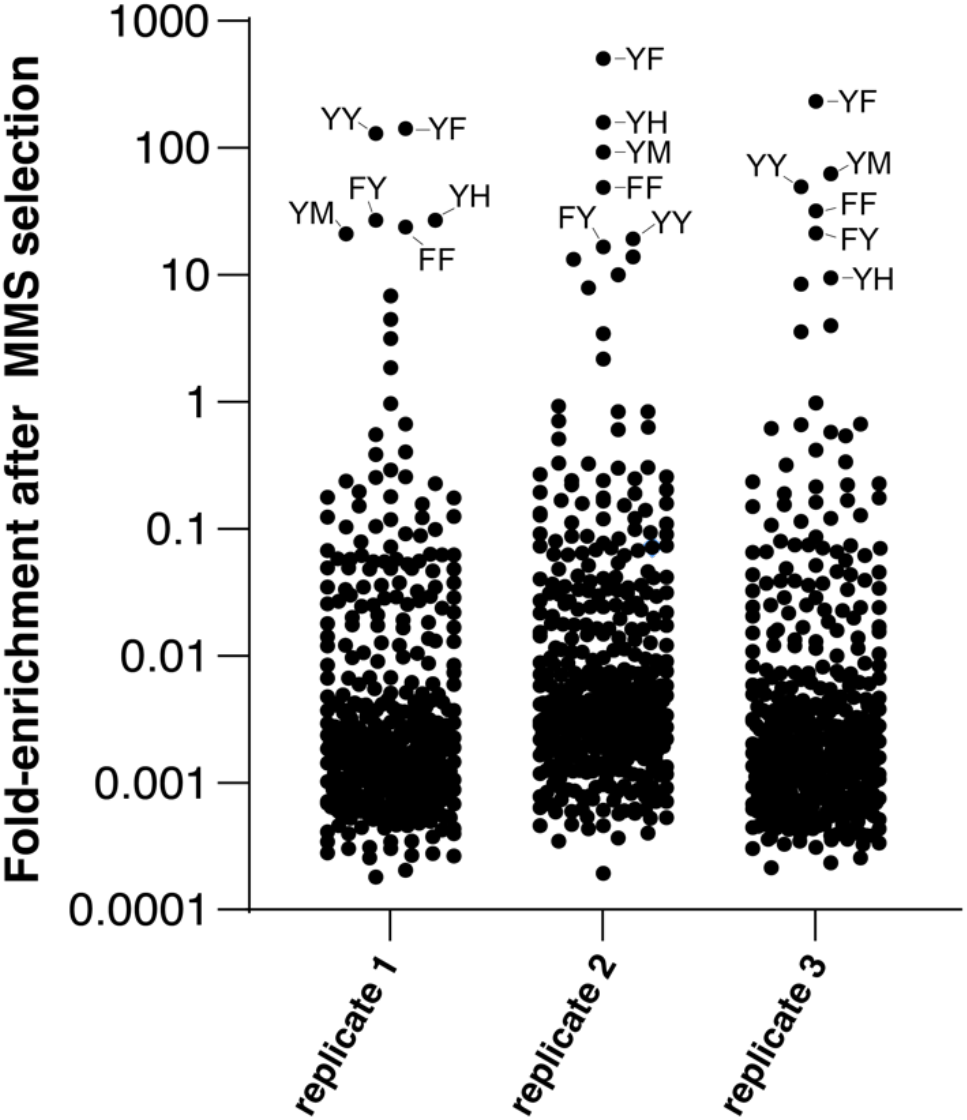
Replicates of high-throughput MMS assays. Scatter plots showing the results of three biological replicates of the high-throughput screen for functional *rad51* variants. The most highly enriched amino acid residue pairs are highlighted.

**Figure S4.**
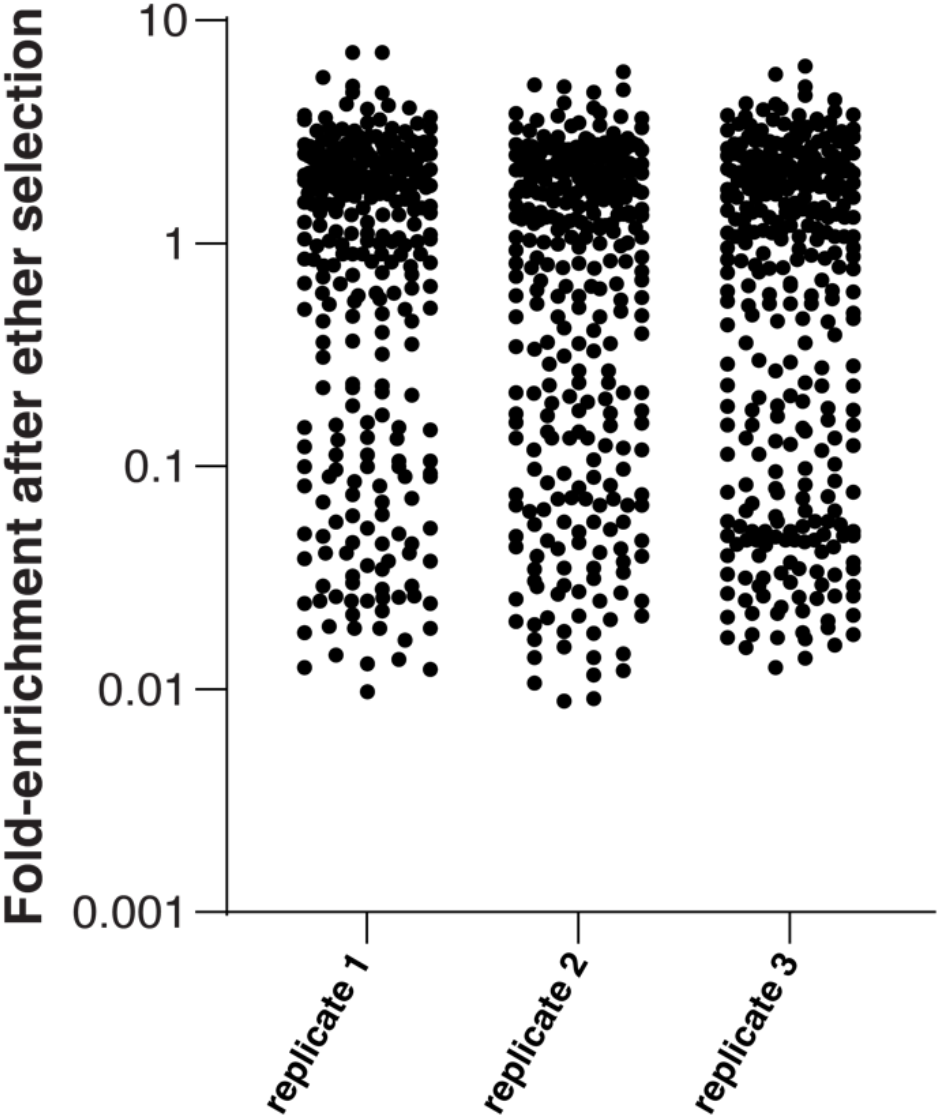
Replicates of high-throughput sporulation assays. Scatter plots showing the results of three biological replicates of the high-throughput screen for functional *dmc1* variants.

**Figure S5.**
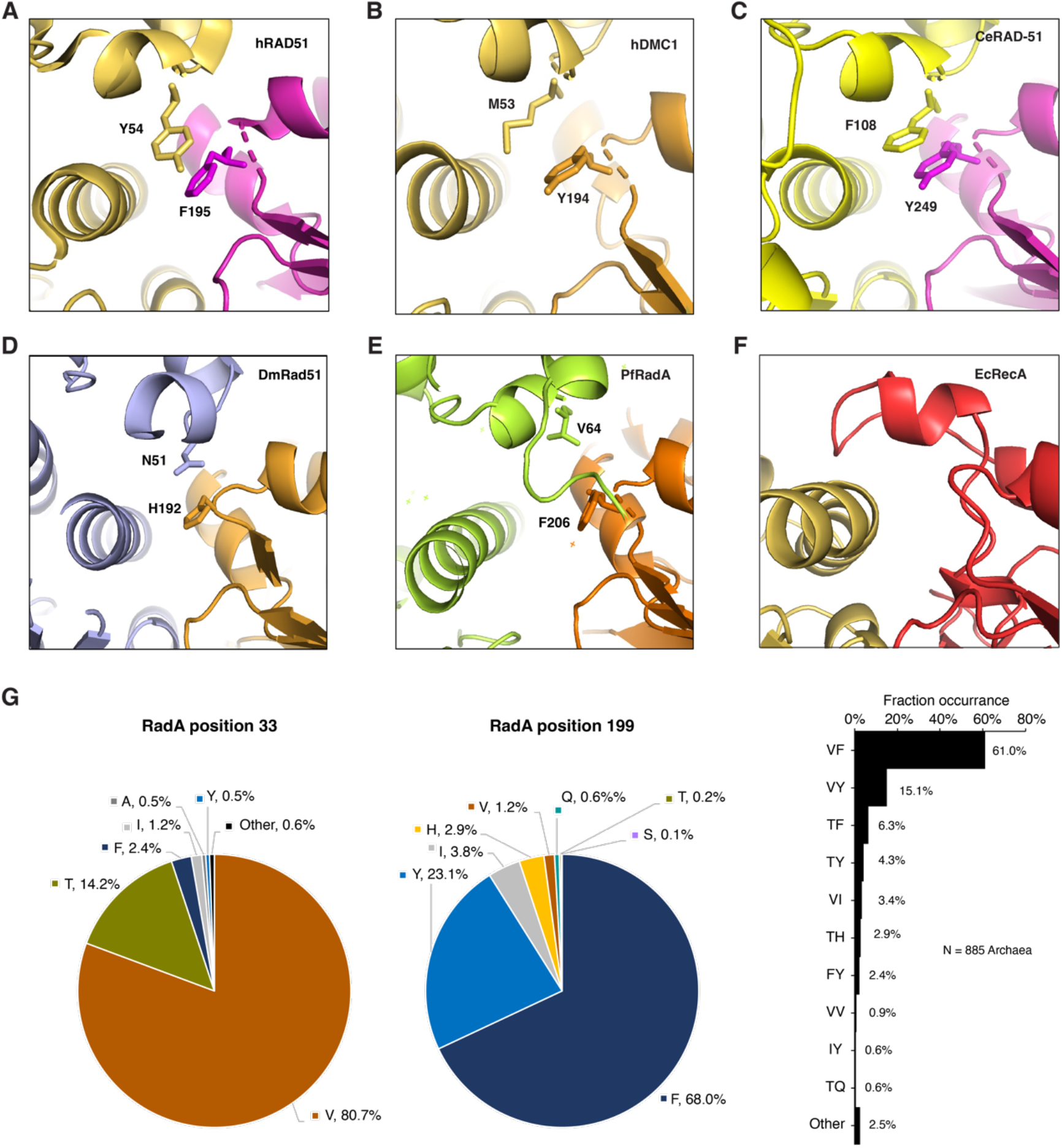
Comparison of interfacial amino acid residues for other members of the Rad51/RecA family. a, Human RAD51 (PDB: 5H1B; (36)). **b,** human DMC1 (PDB: 7C9C; (33)). **c,** *C. elegans* RAD-51 (generated with AlphaFold). **d,** *Drosophila melanogaster* Rad51 (generated with AlphaFold). **e,** *Pyrococus furiosus* RadA (PDB: 1PZN; (85)). **f,** *E. coli* RecA (PDB: 3CMT; (18)). **g,** Interfacial amino acid residue conservation for archaeal RadA. **h,** Naturally occurring RadA interfacial amino acid residue variants.

**Table S1:**
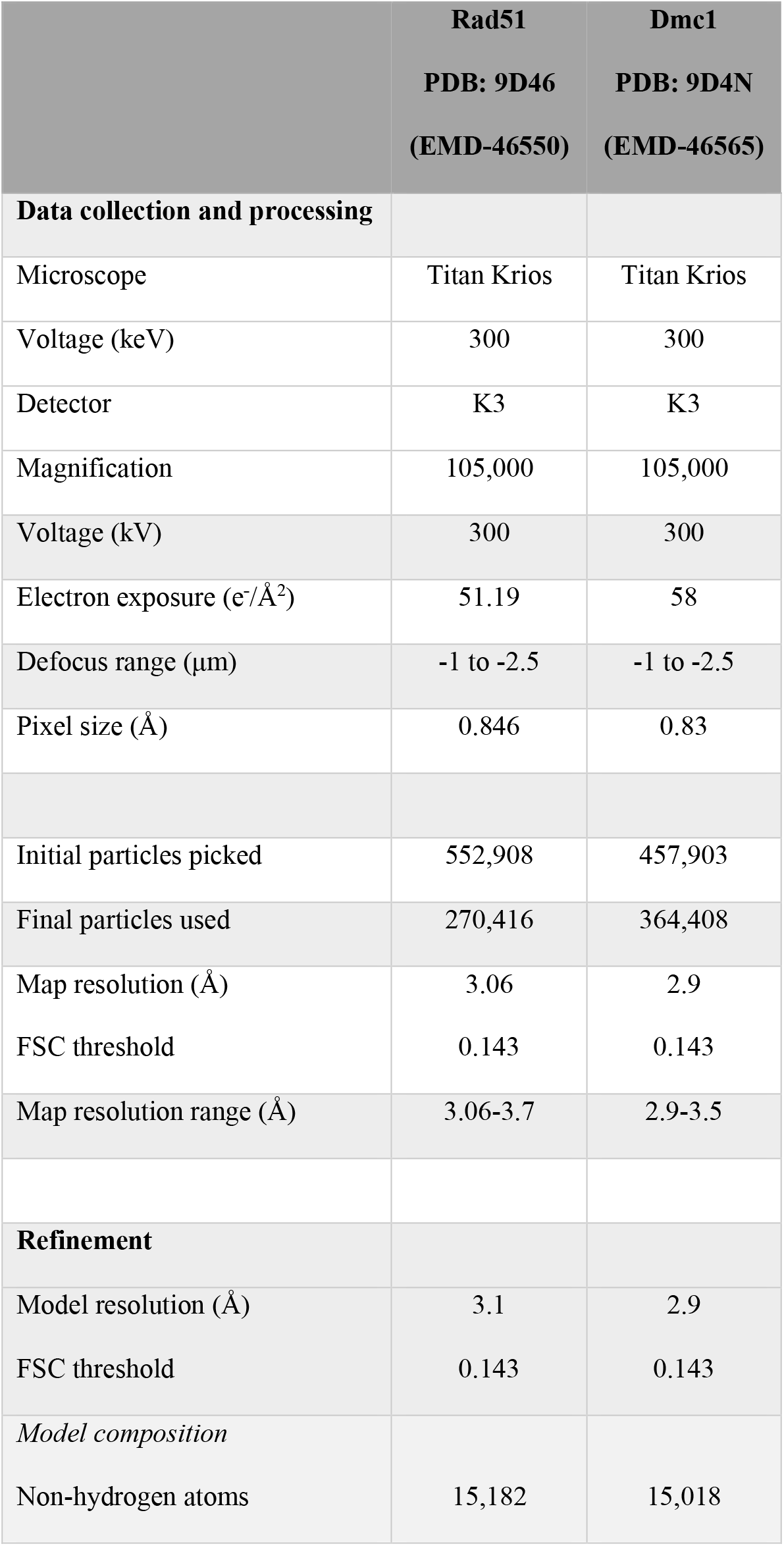
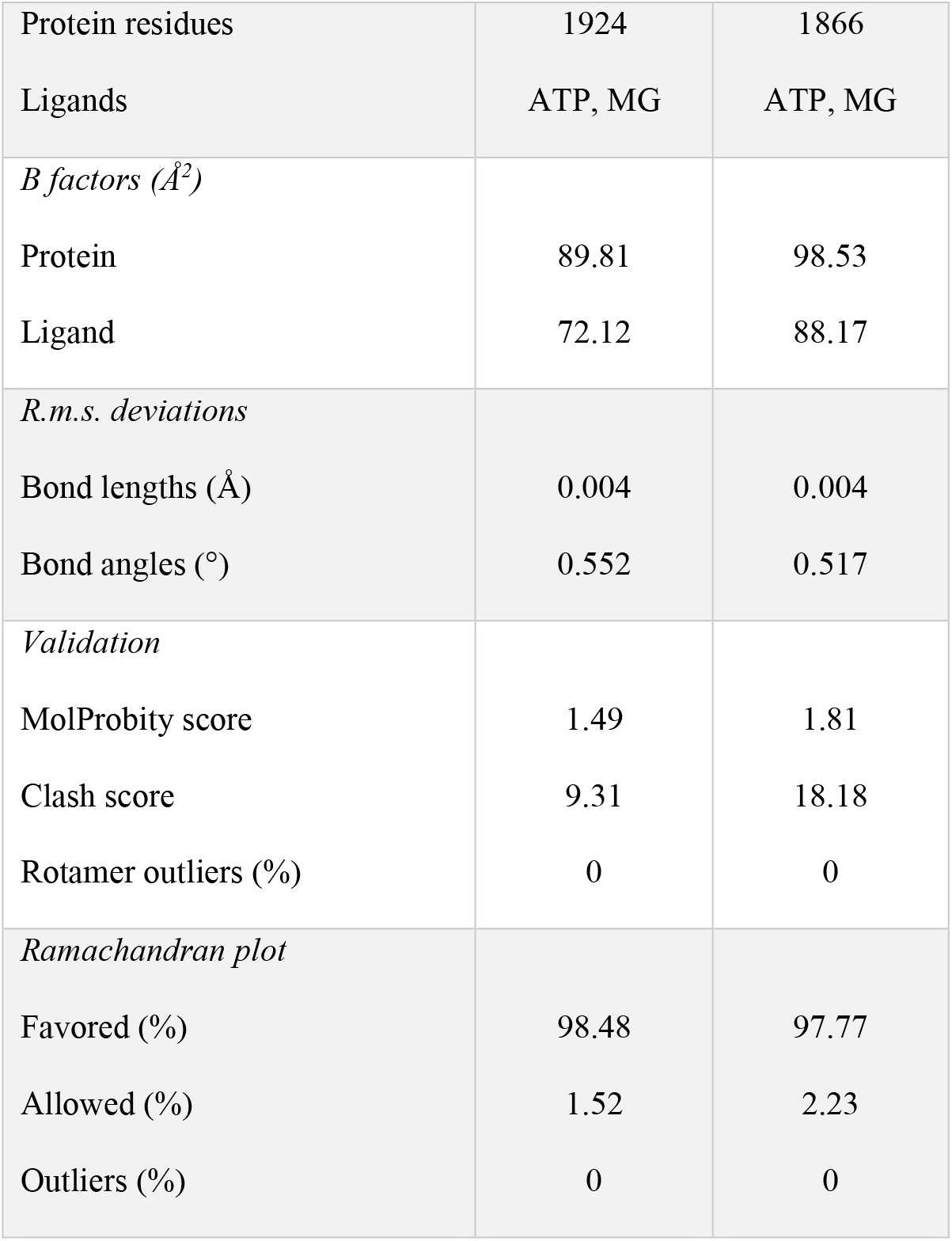

